# Recurrent allopolyploidization events diversify eco-physiological traits in marsh orchids

**DOI:** 10.1101/2021.08.28.458039

**Authors:** Thomas M. Wolfe, Francisco Balao, Emiliano Trucchi, Gert Bachmann, Wenjia Gu, Juliane Baar, Mikael Hedrén, Wolfram Weckwerth, Andrew R. Leitch, Ovidiu Paun

## Abstract

Whole-genome duplication, in particular allopolyploidy, has shaped the evolution of angiosperms and other organisms. Structural reorganization of chromosomes and repatterning of gene expression is frequently observed in early generation allopolyploids, with potential ecological consequences. The relative contributions of environmental and intrinsic drivers to these changes are unknown. We show here that in marsh orchids (*Dactylorhiza*, Orchidaceae), recurrently-formed allopolyploids are characterized by distinct eco-physiologies matching their respective environments, providing us with an excellent study system to address this question. Here we integrate eco-physiological and transcriptomic comparative studies to reveal a complex suite of intertwined, pronounced differences between sibling allopolyploids. We show that *Dactylorhiza majalis* that is distributed in Central and Southern Europe favors meadows with mesic soils. Its sibling allopolyploid *D. traunsteineri* occurs in fens, characterized by soils depleted by macro- and micronutrients, mainly in previously glaciated European areas. We further uncover distinct features in their nutrient transport, leaf elemental chemistry, light-harvesting, photoprotection, and stomata activity, that appear all linked to their distinct ecologies, in particular soil chemistry differences at their native sites. Recurrent polyploidization hence enriches biodiversity through eco-physiological diversification, providing the opportunity for sibling allopolyploids to evolve as distinct evolutionary units, despite pervasive interspecific gene flow.

**Significance Statement:** Whole-genome duplication resulting in polyploidy has underpinned the evolution of flowering plants and other organisms, and is important for many crops. However, the ecological implications of polyploidy remain little understood. Here, we demonstrate that two sibling allopolyploid marsh orchid species prefer distinct habitats, and have evolved a suite of distinctive ecophysiological characters (e.g. nutrient transport, energy harvesting and photoprotection). We argue that the divergence of these characters in the nascent polyploids drove adaptation into distinct ecological niches (low nutrient fens versus meadows with mesic soils), generating ecological barriers that maintains distinct, independent lineages, even in the presence of interspecific gene flow.

## Introduction

Whole-genome duplication is a central force in evolution, and ploidy increase has been estimated to be associated with about one in seven flowering plant speciation events (Wood et al. 2009). In animals, polyploidy is rarer, but multiple cases of currently polyploid insects, fishes, amphibians, and reptiles are known (Otto and Whitton 2000). Moreover, genomic evidence suggests all vertebrates, flowering plants, and some fungi descended from polyploid ancestors, i.e., individuals with three or more sets of chromosomes (Otto 2007; Van de Peer et al. 2017). Polyploidization appears to have significant ecological consequences, as it generally correlates with environmental change or stress (Van de Peer et al. 2017; Novikova et al. 2018), and paleo-polyploidy tends to cluster around past periods of unstable conditions, including mass-extinction events (e.g., Lohaus and Van de Peer 2016).

Polyploidy merges multiple entire genomes in one nucleus, and if polyploids are derived from divergent individuals, or even distinct species (as in allopolyploids) it can trigger a plethora of genomic and transcriptomic responses (e.g., Adams and Wendel 2005). Most neopolyploids show poor fitness and fail to establish (Levin 2002; Husband 2000), but often polyploids are generated with distinctive characters and do establish and become species in their own right.

Certain conditions are considered more favourable for the establishment and spread of polyploids, including the formation of pre-zygotic barriers with diploid parental populations and assortative mating between polyploids (Fowler and Levin 1984; Soltis et al. 2014; Fowler and Levin 2016). These conditions are typically met when allopolyploids are formed. To avoid minority cytotype disadvantage (Husband 2000), the establishment of polyploids appears favoured *i*) in systems that can alter life-history traits, such as flowering time and pollinator specialization, *ii*) when self-incompatibility is absent or breaks-down (Husband et al. 2007; Barringer 2007), *iii*) after long-range dispersal (Felber 1991; Levin 1975; Fowler and Levin 1984; Rodriguez 1996), and *iv*) when the polyploids are able to occupy non-parental niches (Porturas et al. 2019).

Novel ecological properties may arise in early-generation polyploids as a by-product of large-and small-scale adjustments of genome organization and gene regulation that can occur with polyploidy, which with meiotic segregation can lead to transgressive characters, especially in allopolyploids (Paun et al. 2007). Several lines of evidence suggest that polyploids can show novel growth patterns and morphological characteristics compared with their lower ploidy level relatives, with direct implications on complex ecological interactions, even between different trophic levels (Ramsey 2011; Segraves 2017).

Information on early-generation eco-physiological consequences of polyploidy is currently limited (te Beest et al. 2012; Scheriau et al. 2017; Segraves 2017). One ecological consequence of genome multiplication is thought to be an increased nutrient requirement for nitrate and phosphate, necessary for DNA replication and chromatin packing in polyploids compared to diploids, especially in habitats with limited nutrient resources (Ramsey 2011; Glennon et al. 2014; Jeyasingh et al. 2015; Guignard et al. 2016, 2017). System-specific physiological consequences of polyploidy, identified through comparisons between polyploids and related diploids have also been shown, for example, increased salt-tolerance (Chao et al. 2013), alterations in water-use efficiency (Garbutt and Bazzaz 1983), and in photosynthetic activity (e.g., Coate et al. 2012). Such outcomes may allow neopolyploids to range more broadly over the adaptive landscape and access adaptive peaks that would otherwise be unattainable through incremental changes at the diploid level. Early-generation, established polyploids can exhibit larger distributions than their lower ploidy progenitors, especially in habitats that feature less interspecies competition, for example, previously-glaciated regions (Stebbins 1971; Novikova et al. 2018; Rice et al. 2019). Disentangling direct ecological implications of polyploidization in a natural context, relative to adaptation during later evolution, is important for understanding how polyploids spread across niches and their effects on biodiversity (Soltis et al. 2010; Abbott et al. 2013; Jurado et al. 2019). Here, studies of established but young natural polyploids in ecologically-relevant settings are needed.

Recurrence in polyploidization is commonly observed in early polyploid evolution (Soltis and Soltis 1999). Examples include up to 20 independent origins of the allopolyploid *Tragopogon miscellus*, 13 origins for *Draba norvegica*, and 46 for tetraploid *Galax urceola* (Segraves 2017). Recurrent formation of polyploids with different parental genotypes can greatly enhance genetic diversity in the nascent polyploid population (Soltis and Soltis 1999; Soltis et al. 2010). However, independent whole-genome duplications can happen in different ecological contexts too, potentially leading to local adaptation and the formation of polyploid species with different evolutionary trajectories (Jurado et al. 2019), even in the face of gene flow (Novikova et al. 2020).

In this study, we focus on the eco-physiological diversity released by recurrent allopolyploidization events in European marsh orchids (*Dactylorhiza*) in order to identify mechanisms that drive adaptation to different environments and lead to the evolution of distinct allopolyploid species. We investigate two sibling allopolyploids, *Dactylorhiza majalis* and *D. traunsteineri* that are estimated to have formed at different times in the recent half of the Quaternary (Brandrud et al. 2020). They arose from independent, but unidirectional allopolyploidization between *D. fuchsii* (as maternal genome) and *D. incarnata* (Supplementary Fig. S1; Pillon et al. 2007; Hedrén et al. 2008; Balao et al. 2017; Brandrud et al. 2020). Despite their relatively young ages, each of these allotetraploids occupies wide distributions across Europe. *Dactylorhiza majalis* ranges from the Pyrenees to southern Scandinavia, whereas the younger *D. traunsteineri* (Brandrud et al. 2020) has a more disjunct distribution around previously glaciated areas: on the British Isles, northern Europe, and also in the Alps where it grows in sympatry with *D. majalis* (Supplementary Fig. S2A). Both allopolyploids are pollinated through food deceit by inexperienced bumblebees, and such pollination syndrome can promote interspecific cross-pollination (Hedrén and Nordström Olofsson 2018). Indeed, frequent and ongoing gene flow has been documented between the two allotetraploid species in the Alps, but with pervasive traces affecting also populations outside strictly sympatric localities (Hedrén et al. 2008; Balao et al. 2016; Brandrud et al. 2020). A well-documented example of a strongly admixed population is the type locality of *D. traunsteineri* close to Kitzbuhel, Austria, whose plants exhibit the characteristic morphological traits, but generally have a large portion of *D. majalis* alleles (population ALP9 in Balao et al. 2016).

The two sibling allopolyploid species have been suggested to have distinct ecologies, even at localities where the two allopolyploids neighbour each other (Paun et al. 2011). However, no quantitative data has been available on the type and magnitude of ecological differences. *Dactylorhiza majalis* typically grows in wet meadows, whereas *D. traunsteineri* generally prefers marshes and fens with a continuous influx of underground water; both species can benefit from mowing or grazing (Wotavová et al. 2004; Janeckovaa et al. 2006; Sletvold et al. 2011; Djordjević et al. 2016). Furthermore, *D. traunsteineri* is commonly thought to be a poor competitor and tends to grow in sparse vegetation, whereas *D. majalis* generally occupies denser communities where there are more resources (Paun et al. 2010). As established, wide-spread species (Supplementary Fig. S2), these sibling allopolyploids have gone through the filtering effects of natural selection which have acted on their populations since their formation (Pillon et al. 2007).

Using these sibling marsh orchid species, we present a multi-level, integrated investigation of physiology, ecology and gene expression to understand the effects of recurrent allopolyploidization on phenotypic differentiation and adaptation to divergent habitats. Specifically, we quantify the eco-physiological divergence between the two allotetraploids, in particular at the level of macro-climatic conditions, soil chemistry, leaf elemental content, photosynthetic characteristics, and stomatal activity. We further study differential gene expression in a common garden experiment, in order to understand the molecular context that allows the two allopolyploids to occupy distinct niches and to maintain phenotypic distinctiveness, even in the presence of gene flow (Hedrén et al. 2008; Balao et al. 2016; Brandrud et al. 2020).

## Results and Discussion

### The sibling allopolyploid marsh orchids are ecologically distinct

We quantified 18 soil elemental components at 18 localities of the two allopolyploids across Europe (Fig. 1, Supplementary Table S1, Supplementary Fig. S1). Permutation tests found that relative to *D. majalis, D. traunsteineri* prefers soils depleted of several macro- and micronutrients (Fig. 1A, Supplementary Table 2), in particular characterized by extremely low levels of available nitrate (NO_3_^-^), but also phosphate (P) and potassium (K). This indicates that *D. traunsteineri* favours environments with poor mineral nutrition and therefore low competition, for example often occurring alongside carnivorous plants such as sundews (*Drosera* spp; Fig. 1B). In contrast, *D. majalis* prefers richer soils, albeit its populations show in general a wide variance in soil elemental content. Soil pH monitored across several years at 22 localities across the distribution range was also found to be significantly different, with *D. traunsteineri* tending to grow in slightly more alkaline soils than *D. majalis* (Fig. 1A, Supplementary Table S2).

**Figure 1.**
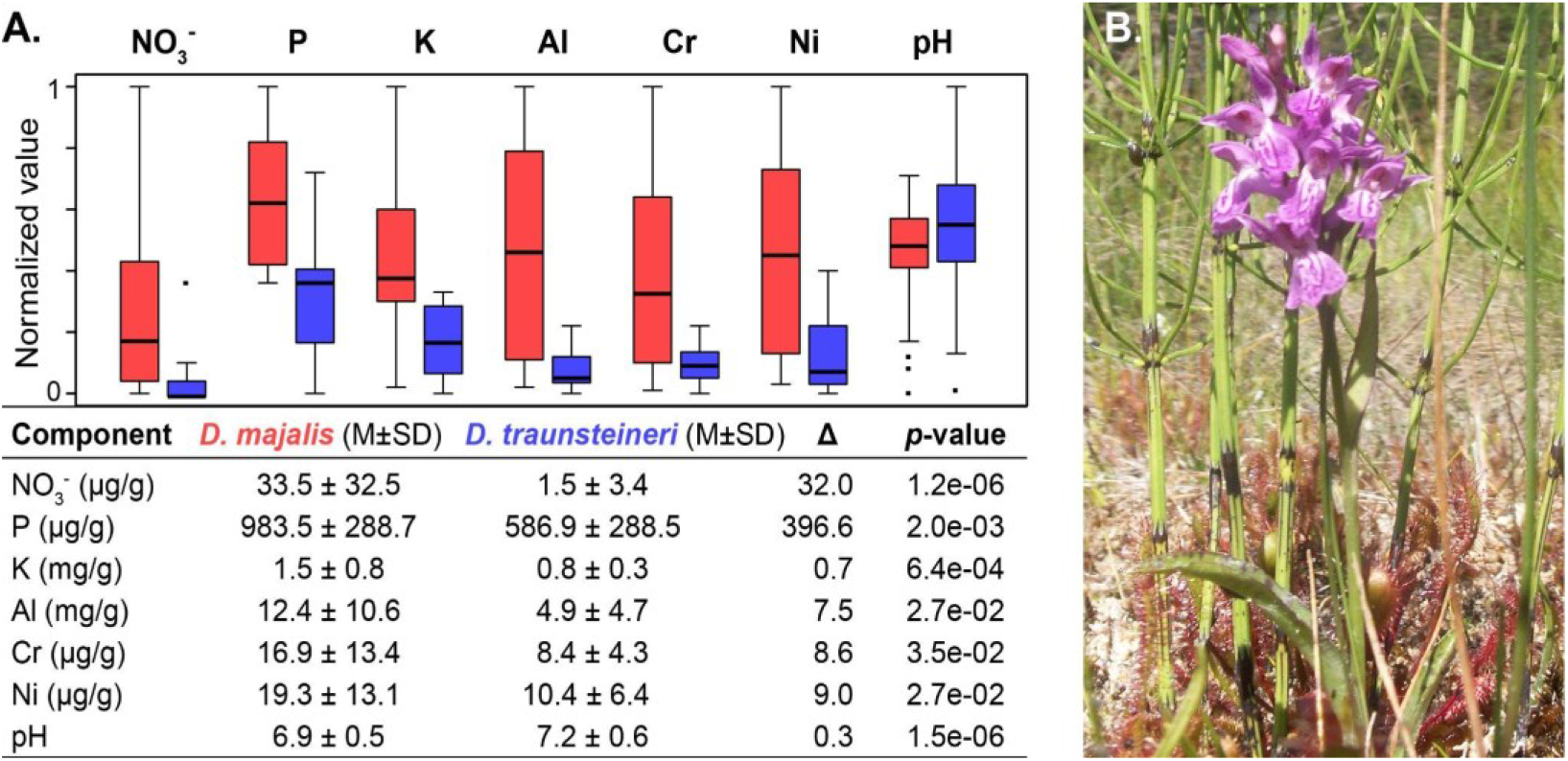
**(A)** Divergent preference for soil chemistry between two sibling *Dactylorhiza* allopolyploids: *D. majalis* (red) and *D. traunsteineri* (blue). Soil elemental profiling for available nitrate (NO_3_^-^), phosphate (P), potassium (K), aluminium (Al), chromium (Cr), nickel (Ni), and soil pH across European populations. The boxplots show normalized values to a 0-1 range using feature scaling, where x_norm_ = (x - x_min_)/(x_max_ - x_min_). The tabular form summarizes the raw data: M, average; SD, standard deviation; Δ, mean difference. The *p*-values are for 1,000 permutation tests. **(B)** *Dactylorhiza traunsteineri* growing alongside carnivorous sundews at a locality on Gotland island, Sweden.

No significant difference was recorded for available ammonium (NH_4_^+^) between the specific soils for the two *Dactylorhiza* allo-tetraploids (Supplementary Table S2). Ammonium can provide a source of nitrogen (N) for plants, particularly in waterlogged soils, but the metabolic processes necessary to assimilate this form of N are different from a nitrate-based metabolism (Xu et al. 2012). In contrast to NO_3_^-^ which is processed in the leaves, NH_4_^+^ is metabolized directly in the roots, a process that requires sugars which have to be transported from leaves to roots. Ammonium uptake can on the other hand affect plant foraging on other nutrients, in particular, K, calcium (Ca), and magnesium (Mg).

To test if the soil elemental profiles specific for each allotetraploid translate to a differential chemistry in the plant, we further analysed leaf chemical content of wild plants at 19 localities across Europe. Elemental analyses (Fig. 2; Supplementary Table S1) showed that relative to *D. majalis*, the narrower leaves of *D. traunsteineri* (Fig. 3A) had significantly lower levels of N and P per weight unit, but not significantly less carbon (C), resulting in higher C:N and C:P ratios. These findings indicate that *D. traunsteineri* has adapted to the low mineral soil composition at its specific sites, optimizing its overall photosynthetic efficiency to fix C in these conditions.

**Figure 2.**
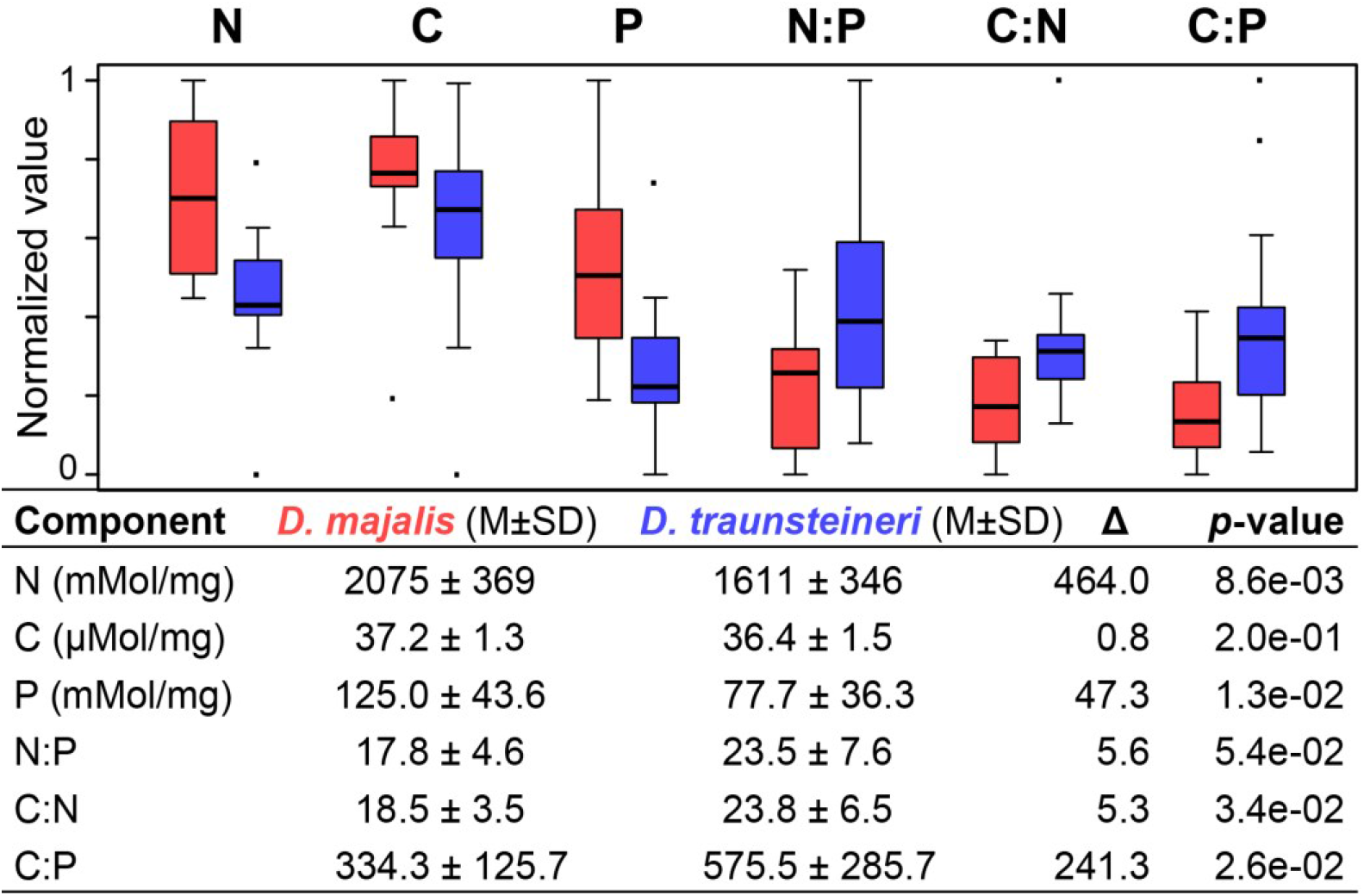
Leaf elemental profiling for wild individuals of *D. majalis* (red) and *D. traunsteineri* (blue) at multiple European localities. Data for nitrogen (N), carbon (C), phosphorus (P) and their pairwise ratios in leaf tissues. The boxplots show normalized values calculated as in Fig. 1. The tabular form summarizes the raw data: M, average; SD, standard deviation; Δ, mean difference. The *p*-values are for 1,000 permutation tests.

**Figure 3.**
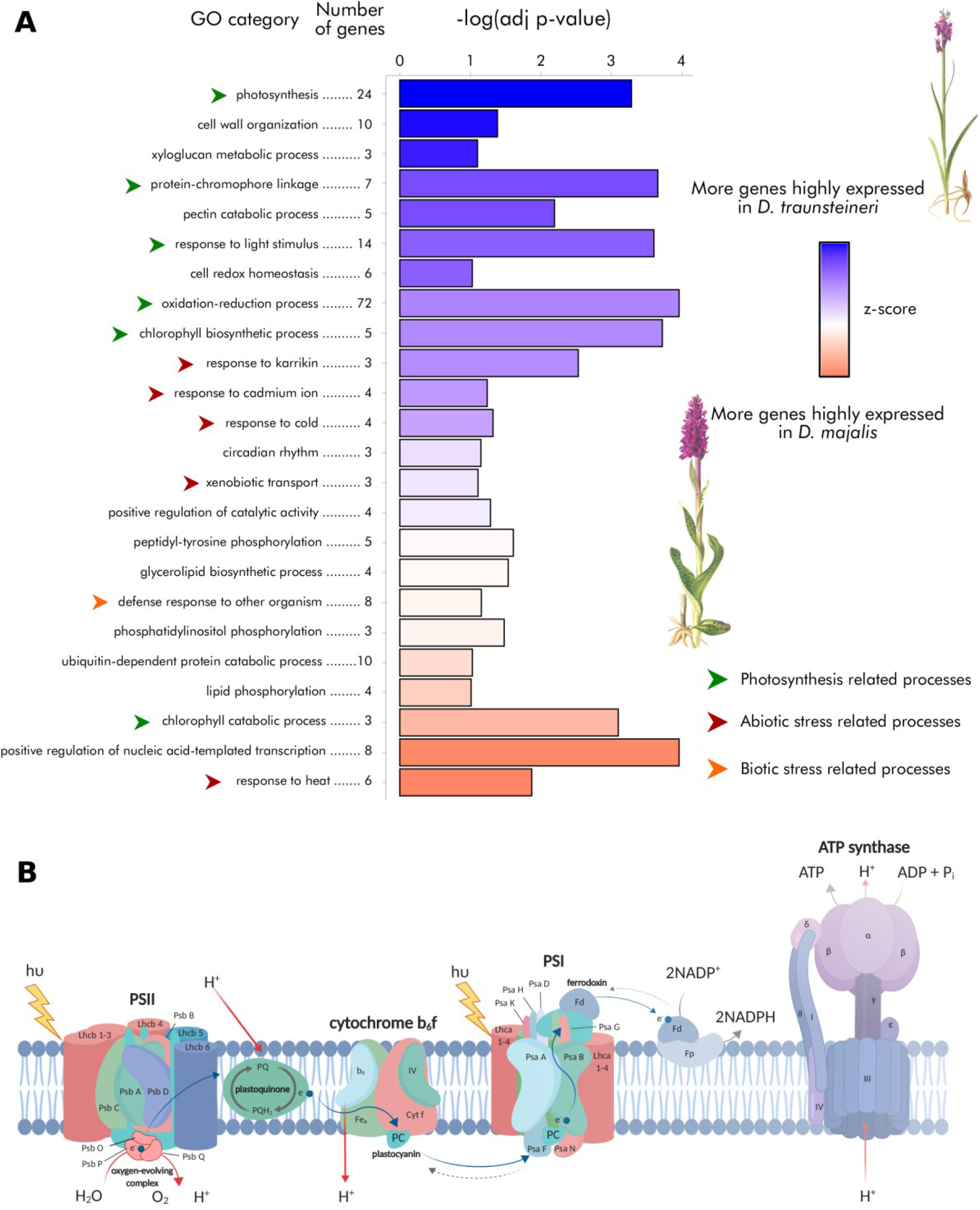
**(A)** Summary of the enriched gene ontology (GO) categories and the number of differentially expressed genes in each of the GO terms. The length of the bars shows the -log(adjusted *p*-value) of the category’s enrichment, and the bar’s colour illustrates the z-score for the category. Blue is for categories where there are more genes higher expressed in *D. traunsteineri*; red is for categories where there are more genes higher expressed in *D. majalis*. Pale colours show that the respective categories have a more balanced ratio of genes overexpressed in either species. Plant illustrations by Erich Nelson (Nelson 1976). **(B)** A summary of photosynthetic reactions drawn with BioRender (https://biorender.com), indicating in red-based colours the components differentially expressed.

We further explored if macro-environmental differences exist between the two allotetraploids, using a selection of bioclimatic and topographic, uncorrelated variables from the CHELSA (Karger et al. 2017) and the ENVIREM databases (Title and Bemmels 2018) at 1 km^2^ resolution for several hundred, manually-curated localities (Supplementary Fig. S2A). Although overlapping, the two climatic niches were evidently not identical (Niche equivalency test, Schoener’s D = 0.32, *p*-value < 0.001; Supplementary Fig. S2B). In particular, temperature seasonality (Bio4), mean temperature of the driest quarter (Bio9), precipitation seasonality (Bio15) and potential evapotranspiration of the driest quarter (PETDriestQuarter) appeared to contribute specificity to the climatic niches of each allotetraploid (Niche divergence test PC1, Student’s t = 16.444, df = 962.31, *p*-value < 0.001; Supplementary Fig. S2B). These results are in agreement with *D. traunsteineri* generally growing in more northern areas (Supplementary Fig. S2A), but also in wetter habitats than its sibling *D. majalis*.

### The preference for distinct ecology is correlated with a complex suite of transcriptomic differences

To examine the functional differentiation between the two sibling allotetraploids, we searched for gene expression differences in a common garden setup. Nineteen representative plants from wild populations of the two allotetraploids across their distribution range (Supplementary Fig. S1B, Supplementary Table S1) had been transplanted and grown in Vienna, Austria for two seasons before RNA isolation, with the aim of removing carry-over environmental differences. The bioinformatics analyses made use of a *de novo* assembled genome for *D. incarnata* v.1.0, newly reported here (Supplementary Information). Principal component analysis (PCA) that summarized the levels of gene expression produced overlapping clusters of the two allopolyploids, and revealed some geographic signal, in particular within *D. traunsteineri*, which likely reflects its disjunct distribution (Supplementary Information, Supplementary Fig. S3). These patterns are in agreement with a PCA constructed based on SNPs derived from the RNA-seq data, and with previous genetic studies (Balao et al. 2016; Brandrud et al. 2020). Both the SNP and gene expression data show in the PCA plots that accessions of *D. traunsteineri* and *D. majalis* from the Alps are intermingled (accessions shown with squares in Supplementary Fig. S3), confirming hybridization in sympatry. Yet despite on-going hybridization, the phenotypic differences between *D. majalis* and *D. traunsteineri* are stable and distinctive, even for individuals of the two allotetraploids growing in close proximity. In addition, the sympatric populations essentially remain binary from the point of view of ecological conditions (Paun et al. 2010; Paun et al. 2011; Balao et al. 2016).

Differential gene expression tests revealed that 316 genes (2.5% of those retained after count-per-million based filtering) were significantly overexpressed in *D. majalis*, and 343 (2.7% of retained genes) significantly over-expressed in *D. traunsteineri* (Fig. 3A; Supplementary Fig. S4). Photosynthesis (GO: 0015979) and related processes were by far the most affected by differential expression between *D. majalis* and *D. traunsteineri* (Supplementary Table S3). In particular, relative to *D. majalis, D. traunsteineri* shows an increased expression of the Lhcb1-Lhcb4 antenna proteins that are part of the light-harvesting complex (LHC), which captures and delivers excitation to photosystems (Fig. 3B; Li et al. 2004). The LHC overexpression in *D. traunsteineri* may be linked to a lower relative chlorophyll content compared to *D. majalis* (see below). Also, overexpressed in *D. traunsteineri* are several components of the oxygen-evolving complex of the photosystem II (PSII; Fig. 3B) that are responsible for catalysing the cleavage of water to oxygen, protons and electrons (Raymond and Blankenship 2004). *Dactylorhiza traunsteineri* has also higher expression in the Photosystem I (PSI) reaction centres II (psaD), III (psaF), V (psaG), XI (psaL), N (psaN), and O (psaO) (Fig. 3B), which are mediating the primary function of the PSI (i.e., electron transfer from plastocyanin to ferredoxin; Jensen et al. 2004). Notably, 10.9% of the differentially expressed genes (i.e., mostly over-expressed in *D. traunsteineri*) are involved in oxidation-reduction, a process that also shows one of the strongest over-representations in the enrichment test (Fig. 3A). Finally, also related to photosynthesis, genes linked to lightharvesting in PSI, chlorophyll biosynthesis processes, response to blue light, and response to light stimulus are overexpressed in *D. traunsteineri* compared to *D. majalis* (Fig. 3A).

### The sibling allopolyploid marsh orchids are physiologically distinct

To complement the transcriptomic and ecological results presented above, and confirm the photosynthetic differences recovered via RNA-seq, we phenotyped multiple accessions of both allopolyploids in the wild for leaf photosynthetic parameters using a MultispeQ v.1.0 (Kuhlgert et al. 2016). For a total of 38 replicated individuals from three Alpine populations of *D. traunsteineri* and 20 individuals from two Alpine populations of *D. majalis* we quantified photosynthetic parameters (parameters defined in Fig. 4, Supplementary Table S4).

**Figure 4.**
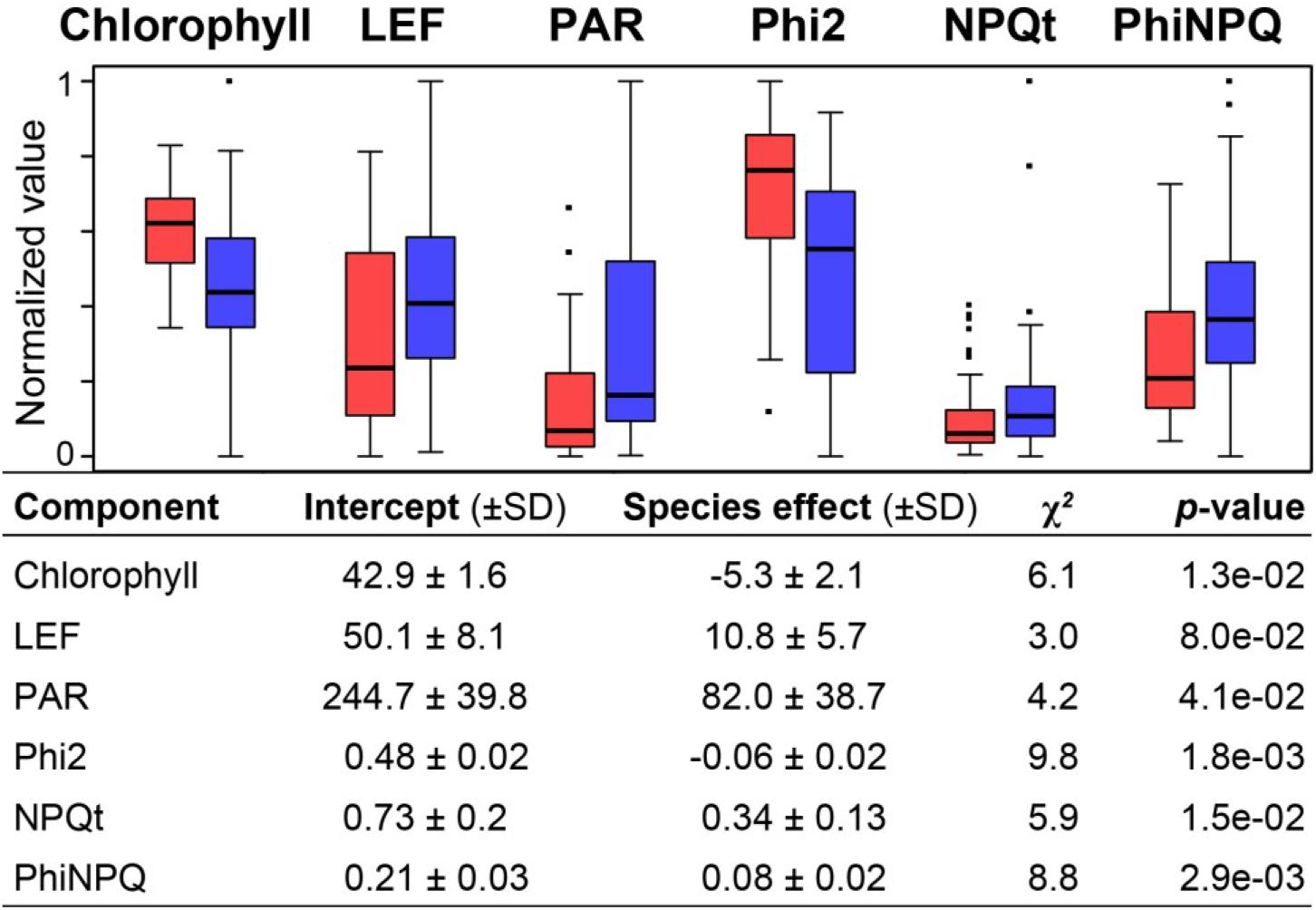
Photosynthetic characteristics for the sibling allotetraploids *D. majalis* (red) and *D. traunsteineri* (blue) in the Alps. LEF, Linear Electron Flow; PAR, photosynthetically active radiation (light intensity); Phi2, Quantum yield of Photosystem II; NPQt, non-photochemical quenching; PhiNPQ, ratio of incoming light that goes towards non-photochemical quenching. The boxplots show normalized values calculated as in Fig. 1. The tabular form gives the details of the likelihood ratio between the null model with time, date and individual measurements as random variables, and the full model with species added to the model as a fixed variable. SD, standard deviation.

Compared to *D. majalis*, the narrower leaves of *D. traunsteineri* showed significantly less relative chlorophyll content, which in effect is a measure of leaf “greenness” (Fig. 4). This could reflect the specific nutrient poor soils preferred by *D. traunsteineri* and the nitrogen deficiency quantified in its leaves, as nitrogen is required for chlorophyll synthesis. It is of relevance here to note that we had observed in the RNA-seq results in common garden conditions overexpression of light harvesting proteins (Fig. 3) of *D. traunsteineri* compared to *D. majalis*. As *D. traunsteineri* prefers poor soils (Fig. 1A), it is likely that chlorophyll deficiency became ameliorated by an increased activity of light harvesting proteins. Such alterations to light harvesting complexes may make the regulation of captured light more difficult. In the wild, *D. traunsteineri* used 82.0 µmol photons∗ *m*^−2^ ∗ *s*^−1^ ± 38.7 more incoming light (400 nm to 700 nm) to drive photosynthesis (PAR), compared to *D. majalis*. However, the quantum yield (Phi2), i.e., the number of excited electrons that go into the Photosystem II, was significantly lower in *D. traunsteineri* than in *D. majalis*. Altogether, the amount of incoming light regulated away from photosynthetic processes in order to reduce damage to the plant (i.e., non-photochemical quenching, phiNPQ) was also significantly higher in *D. traunsteineri* (Fig. 4). The increased phiNPQ in *D. traunsteineri* compared to *D. majalis* is also likely reflected in the overexpression of oxidation-reduction processes (strongly enriched in differential expression tests; Fig. 3A), with a role in coping with residual oxygen radicals. This suggests that *D. traunsteineri* needs to implement a stronger photoprotective strategy to regulate its photosynthetic activity via quenching, potentially due to the limiting amount of chlorophyll, itself likely reflecting a deficiency of nitrate and other nutrients at those respective sites (Demmig-Adams and Adams 2006; Lopez-Jurado et al. 2020).

Finally, we measured leaf stomatal conductance for water and net CO_2_ exchange in a controlled lab environment using a portable gas exchange fluorescence system GFS-3000 (Heinz Walz, Effeltrich, Germany). We found that both species maintained their stomata open during night-time, and hence gas exchange through stomata occurred throughout the night (Fig. 5), likely using evapotranspiration to maintain xylem transport to accommodate for nutrient scarcity. During the night, *D. majalis* showed a negative net CO_2_ (i.e., a release of CO_2_) in contrast to *D. traunsteineri*. In addition, during daytime *D. traunsteineri* transpires ca 20% less water, and this higher water-usage efficiency complements a ca 50% higher CO_2_ net assimilation rate compared to *D. majalis*. What this means for the enzyme apparatus and the light to energy conversion of the species needs to be elucidated in future research. As the transcriptomic aspects and gas exchange results have been obtained in a common garden, it appears likely that these physiological features are genetically encoded.

**Figure 5.**
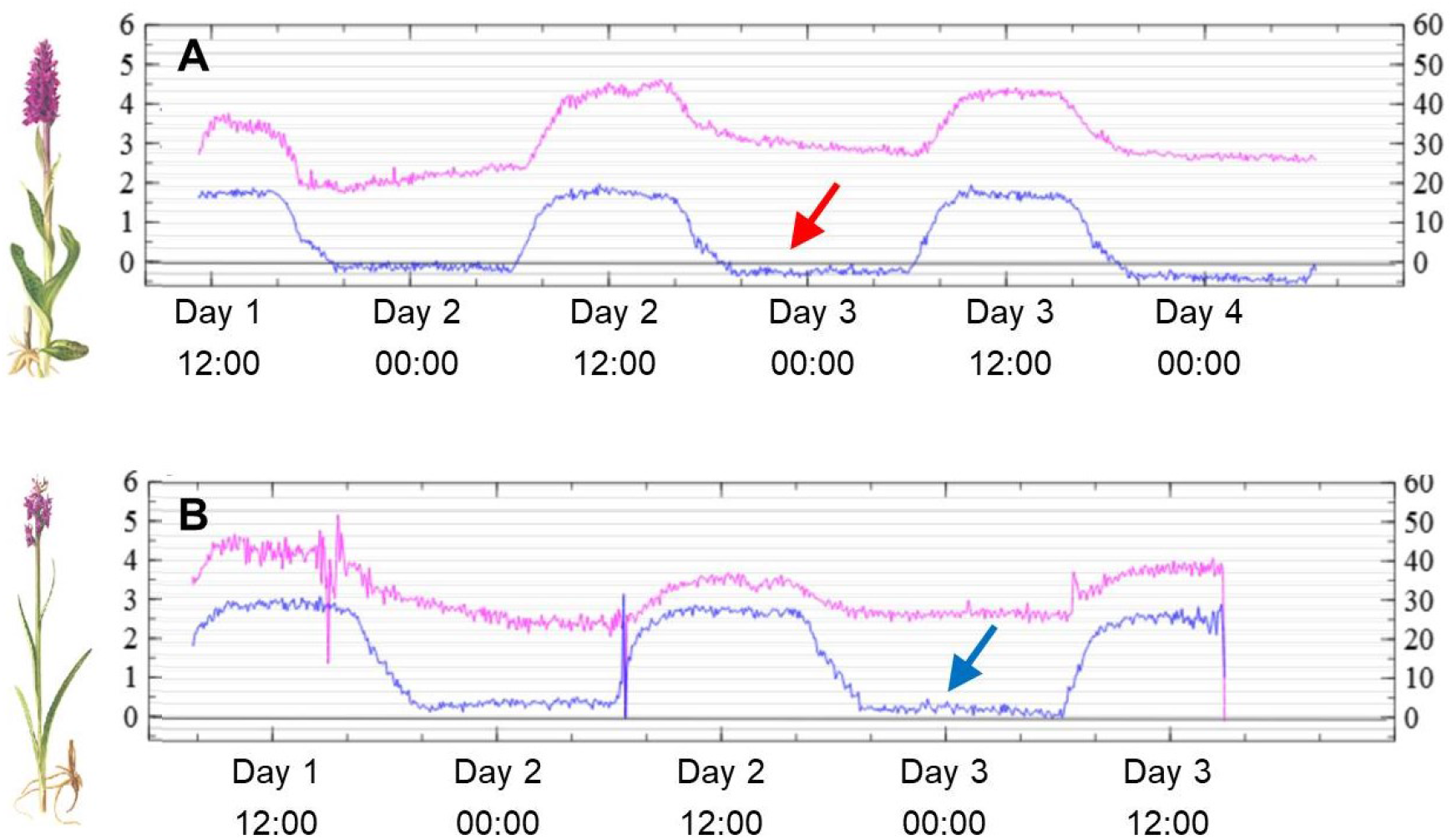
Examples of leaf stomatal conductance for water vapour (pink line, measured in mmol m^-2^ s^-1^ on the right-side Y-axes) and net CO_2_ assimilation rate (blue line, measured in umoles m^-2^ s^-1^ on the left-side Y-axes) for an individual of *D. majalis* **(A)** and one of *D. traunsteineri* **(B)**. The levels of stomatal conductance and CO_2_ exchange fluctuate with a twelve-hour day/night cycle. Stomatal conductance is lower in *D. traunsteineri* which shows that its leaves do not open stomata as much as *D. majalis*, resulting in less water loss (higher water use efficiency) for nutrient acquisition via xylem. At night, the net CO_2_ exchange is slightly negative for *D. majalis* (red arrow) meaning that it releases more CO_2_ through respiration during nighttime, in contrast to *D. traunsteineri* (blue arrow), which also features 50% higher CO_2_ assimilation rate during daytime. Plant illustrations by Erich Nelson (Nelson 1976).

### Conclusions

Although *D. majalis* and *D. traunsteineri* arose from similar, unidirectional allopolyploidization events (Brandrud et al. 2020), we uncovered major and intertwined environmental, physiological and transcriptomic differences between them. Despite the frequent gene flow between the two allotetraploids (Hedrén et al. 2008; Balao et al. 2016; Brandrud et al. 2020), this appears not to homogenize the phenotypes of the allopolyploids. This suggests the existence of strong ecological divergent selection that acts at particular loci, and is responsible for maintaining distinct allopolyploid species. Their distinct ecological preference has arisen from independently-derived allopolyploidy events involving the same diploid progenitor species. The strength of the ecological segregation seems to be sufficient to maintain species integrity in the face of gene flow. This conclusion supports the evidence that ecological differentiation is a major driver behind orchid diversification in general (e.g., Tupac Otero and Flanagan 2006; Ackermann et al. 2007).

The pace of divergence of allopolyploids from the initial, neopolyploid genomic background will depend heavily on population parameters such as gene flow, generation time and mating systems (Yang et al. 2018), or phenological parameters such as the number of flowers per individual and the complexity of pollinator interactions. Given their widespread distribution, possibilities for long-range dispersal and the success of both polyploids, we assume genetic drift isolating these species is likely to have been a minor force for these established allopolyploids in the recent past and the present. However, when populations were smaller, around the time of their formation, stochastic processes have likely been more prominent, and established variation that enabled the colonization of different habitats.

It remains unclear whether the eco-physiological differences between the allotetraploids are due mainly to selection on new polymorphisms, selection on standing variation from within the parental sub-genomes, or drift during early stages after allopolyploidization. Nevertheless, we show in this study that establishment of recurrent allopolyploids into distinct niches can happen relatively rapidly (te Beest et al. 2012) and this can significantly contribute to maintaining distinct phenotype and evolutionary trajectories, even when sharing the same ploidy, a similar genetic background and in the face of a pervasive gene flow.

## Materials and methods

### Soil and leaf elemental analyses

Soil samples were collected for nine localities of *D. majalis* (five from the Alps, two from the Pyrenees and two from Scandinavia) and nine localities of *D. traunsteineri* (three from the Alps, three from the Britain and three from Scandinavia; Supplementary Table S1). Two composite soil samples, each including three subsamples in approximately equal proportion, were profiled per locality according to Supplementary Information. Significance of distribution differences for soil characteristics between the two allotetraploids were tested using permutation tests with the *permTS* function in the R package ‘perm’ (Fay and Shaw 2010).

For leaf elemental measurements plant tissue was collected at nine *D. majalis* localities (five from the Alps, two from the Pyrenees and two from Scandinavia and) and at ten *D. traunsteineri* populations (three from the Alps, five from Britain and two from Scandinavia; Supplementary Table S1), stored in silica gel, and analysed according to Supplementary Information. Significance was assessed using permutation tests as explained above.

### Macro-environmental climatic niche characterization

Occurrence information for both allotetraploid species was collected from Global Biodiversity Information Facility (GBIF; https://www.gbif.org) and were further manually curated, retaining 298 localities for *D. majalis* and 393 localities for *D. traunsteineri*. The analyses followed Balao et al. (2017), except that the environmental data for these localities were extracted from the CHELSA (Karger et al. 2017) and ENVIREM databases (Title and Bemmels 2018) (see Supplementary Information).

### RNA-seq analyses

We collected eleven adult *D. traunsteineri* plants from nine localities and eight adult *D. majalis* plants from eight localities throughout Europe and grew them in a common garden setting for two years in Vienna, Austria (see Supplementary Fig. S1, Supplementary Table 1). Every year, these orchids store nutrients into a new tuber to support growth in the following year. Leaf tissues were fixed in RNAlater for all accessions on the same day and in a similar developmental stage. The wet lab procedure followed the details given in Balao et al. (2017). The RNA-seq libraries were sequenced as directional, 100bp paired-end reads with Illumina HiSeq. Reference genome assembly, RNA-seq read quality control, mapping, normalization and differential expression analyses are described in Supplementary Information.

### Photosynthesis and gas exchange

Photosynthesis was measured with two replicate measures on different leaves for each accession in the field. The measurements were taken at two localities in the Alps, where *D. majalis* and *D. traunsteineri* are within a few hundred meters of each other (i.e., ALP9, ALP13; Supplementary Table 1) and one population where only *D. traunsteineri* grows (ALP8). We analysed the data with mixed linear models according to Supplementary Information. Despite the proximity of the two species at these sympatric localities and our randomized sampling, the MultispeQ measurements uncovered a difference in the ambient conditions around the two allopolyploids. In accordance with its wet and exposed habitats (i.e., low surrounding vegetation) *D. traunsteineri* experienced on average 3.5 % ± 1.1 (std. error) more ambient humidity (*p* = 0.0045) and a 1.4 ± 0.4 °C higher temperature (*p* = 2e-10) than *D. majalis*.

Finally, we measured leaf stomatal conductance for water and net CO_2_ exchange in a controlled lab environment set to 25 °C during day and 23 °C in the night, 50% air humidity, 12 h photoperiod at 400 umoles m^-2^ s^-1^ with a LED simulated sunlight spectrum set to 5,800K, watering the pots daily to field capacity.

## Data accessibility

The raw Illumina sequencing data for *D. majalis* and *D. traunsteineri* are deposited on NCBI SRA (PRJNAX). The new *D. incarnata* genome assembly is available from GenBank (accession number TBC). The table of counts and scripts are available on GitHub (https://github.com/twolfe/dactylorhiza).

## Acknowledgments

We would like to thank Alexander Athanasiadis, Richard Bateman, Connor Bottomley, Mark Chase, Laure Cyveyrel, Marcel Hirsch, Christian Lexer, Maria Teresa Lorenzo, Martin Pontz, David Pressler, Judith Trunschke and Claus Vogel for their contribution to the results presented here. Faculty members, the students and the scientific advisory board of the Vienna Graduate School of Population Genetics (www.popgen-vienna.at/) are acknowledged for numerous discussions and feedback on this work. Sequencing was performed at the Vienna BioCenter Core Facilities (VBCF; https://www.viennabiocenter.org/). Computational resources were provided by the Vienna Scientific Cluster (VSC) and the Life Science Compute Cluster (LiSC) of the University of Vienna. Natural England and the Forestry Commission (UK), and regional county administrations in Austria, France, and Sweden are acknowledged for issuing necessary collecting permits. This research was funded by the Austrian Science Fund (FWF) through the START grant Y661-B16 to O.P., and the doctoral programme grant W1225-B20 to a faculty team including O.P.

## Author Contributions

Study conceived and designed by OP and TMW. Laboratory work conducted by TMW, GB, WG, JB. Bioinformatics and statistical analyses conducted by TMW and FB. Interpretation of the results was undertaken by TMW, FB, ET, GB, MH, WW, AL and OP. The manuscript was drafted by TMW and OP, and was revised and approved by all authors.

## Notes

### Competing Interest Statement

The authors have declared no competing interest.

