## Supplementary Information for "Recurrent allopolyploidization events diversify eco-physiological traits in marsh orchids"

#### Soil and leaf elemental analyses

The soil samples were collected in spring 2017. The soil was dried and the elemental composition was measured at facilities of the University of Natural Resources and Life Sciences, Vienna, Austria using a Truspec CN Elementalanalyser (LECO Corporation). Available nitrate and ammonium were extracted in 1M KCl and measured on a plate reader photometer (Bio-Rad). A Direct Soil pH Meter HI99121 (Hanna Instruments) was used to measure soil pH at *ca*. 7 cm depth in the close proximity of plants at 22 European localities in multiple years from 2010 to 2015 (Supplementary Table S1).

To measure N and C in the leaf tissue, 1-2 mg oven-dried ground sample was mixed with a urea solution and the mixtures were quantified on a Mass Spectrometer (Integra 2, Sercon, UK). One blank and two case-Rs as reference were also loaded for every ten samples. To measure P, 0.8-2 mg oven-dried, ground tissue was mixed with 0.15 g potassium persulphate and 1 ml 1N sulphuric acid and extracted at 121 °C for 40 min. The P quantification was finally performed on a Skalar machine (Segment Flow Analyser, San++, Skalar Analytical B.V., Breda, The Netherlands).

#### Macro-enviromental parameters

A total of 958 *D. majalis* and 631 *D. traunsteineri* localities (Supplementary Fig. S2A) were extracted from the Global Biodiversity Information Facility (GBIF) using the package 'rgbif' (Chamberlain et al. 2017) in R software v.3.4.2. Localities were manually-curated based on expert knowledge, duplicates were removed, and locally dense sampling was reduced by thinning the records to one per 10 $km^2$ grid cell size. We extracted 36 environmental variables with relevance to the plant eco-physiological conditions from the CHELSA database (Karger et al. 2017) and the ENVIREM database (Title and Bemmels 2018) at ~1 $km^2$ resolution. We used the variance inflation factor (VIF; threshold of 10) to remove highly correlated environmental variables in the dataset. This left only 15 of the 36 environmental variables, summarizing isothermality (BIO3), temperature seasonality (BIO4), mean temperature of the wettest quarter (BIO8), mean temperature of the driest quarter (BIO9), and precipitation seasonality (BIO15), precipitation of warmest quarter (BIO18), precipitation of coldest quarter (BIO19), index of the degree of water deficit below water need (aridityIndex), sum of mean monthly temperature for months with mean temperature greater than 5℃ multiplied by number of days (growingDegDays5), minimum temperature of the warmest month (minTempWarmest), mean monthly Potential Evapotranspiration (PET) of the coldest quarter (PETColdestQuarter), mean monthly PET of the driest quarter (PETDriestQuarter), monthly variability in PET (PETseasonality), mean monthly PET of the wettest quarter (PETWettestQuarter) and topographic wetness index (topoWet).

To compare the macro-environmental niche of the sibling allotetraploids, we calculated the kernel smoothed densities of each occurrence data along environmental axes from a Principal Component Analysis (PCA-env; Broennimann et al. 2012). For background, we extracted environmental data from 3,000 spatially random localities within a buffer of 100 km around the occurrence localities. Then, we estimated the environmental niche overlap and performed the niche equivalence test (Broennimann et al. 2012) comparing the observed overlap (Schoener's D-statistic) with a null distribution based on 1,000 replicates and an environmental grid resolution of 500 x 500 pixels in the R package *ecospat* v. 3.2. (Di Cola et al. 2017). In the niche divergence test, we used Student's t-tests to evaluate the differences in PCA scores for both allotetraploids.

### Reference genome assembly and annotation

We assembled *de novo* a draft genome for a diploid individual of *D. incarnata* from Austria (48° 11.6' N, 16° 29.0' E) using 177.6 Gb PacBio long reads (i.e., ca. 50.7× coverage). PacBio library preparation and sequencing of 18 SMRT cells on a Sequel I instrument was performed at the sequencing facility of the Vienna BioCenter Core Facilities (VBCF; https://www.viennabiocenter.org/). The assembly was performed with Canu v.1.8 (Koren *et al*. 2017) with default settings, including the recommended correctedErrorRate 0.045. Based on the PacBio reads, the raw assembly was scaffolded with SSPACE-LongRead v.1-1 (Boetzer and Pirovano 2014) and polished with Arrow v.2.3.3 (available from https://github.com/PacificBiosciences/GenomicConsensus). The draft *D. incarnata* genome v.1.0 contains 3,418 scaffolds and has an N50 size of 7.4 Mb, recovering a total length of 3.27 Gb. This corresponds to 94.3% of the estimated genome size (1C = 3.55 pg, Aagaard *et al*. 2005).

The genome was structurally annotated *ab initio* using Augustus (Stanke *et al.* 2006) and GeneMark-ET (Lomsadze *et al*. 2014), as implemented in BRAKER1 v.2.1.0 (Hoff *et al*. 2016) with the options --softmasking=1 --filterOutShort. Mapped mRNA-seq data of the same accession was used to improve *de novo* gene finding. The annotation was further improved using MAKER-P v.2.31.10 (Campbell et al. 2014) supplying gene models identified using BRAKER1. Before MAKER analyses, we generated a custom repeat library for repeat masking using RepeatModeler v.1.0.11 (http://www.repeatmasker.org/RepeatModeler/). This process masked as repeat sequences 83.7% of the genome reference. A transcriptome was assembled using Trinity v.2.4.0 (Haas *et al*. 2013) based on the mRNA-seq data, and this has been used in MAKER-P as expressed sequence tags (EST). The annotation used as additional evidence (declared as *altest*) the transcriptome of the related *Orchis italica* (De Paolo *et al*. 2014) and protein sequences of *Phalaenopsis equestris* (Cai et al. 2015). Structural annotations identified a total of 52,665 protein-coding genes. The gene models were functionally annotated using Blast2GO v.5.2.5 (Götz *et al*. 2008), recovering full annotations for 31,394 genes. Finally, we evaluated the completeness of the genome assembly by searching our gene models against the BUSCO v.3 Liliopsida odb10 dataset (Simão *et al*. 2015). A total of 89.2% of the set of single-copy conserved BUSCO genes were found within our annotated genes, out of which 15.3% genes were recorded as duplicated.

### RNA isolation, library preparation and PCA analyses

Leaf tissue was collected in RNAlater in the morning of 14th May 2014, and was left at 4 °C overnight, before being transferred to -80 °C for storage. Total RNA was extracted using the RNeasy Plant Mini Kit (Qiagen) following the manufacturer's instructions. The purified RNA was stored at -80 °C. The concentration of RNA extracts was first measured with a NanoDrop ND-1000 Spectrophotometer (Thermo Scientific) and its purity was estimated according to the wavelength ratio of A260/280. The quantification and quality of the RNA were then confirmed with an RNA 6000 Nano-kit on a 2100 BioAnalyzer (Agilent Technologies). The ribosomal RNA was depleted with the RiboMinus$^{TM}$ Plant Kit for RNA-Seq (Invitrogen) following the manufacturer's protocol. RNA fragmentation was done by hydrolysis for 2 min at 94 °C and used 2 µl 10x buffer RT (SuperScript III First-Strand Synthesis System for RT-PCR, Invitrogen) plus 75mM $MgCl_2$ in 12 µl concentrated RNA. After a cleaning step with an RNeasy spin column (from an RNeasy Plant Mini Kit, Qiagen), the first cDNA strand has been synthesized with the Superscript III First-Strand Synthesis System for RT-PCR (Invitrogen) and random hexamers. Surplus dNTPs have been eliminated with a Mini Quick Spin Column for DNA (Roche). The second strand cDNA synthesis has been performed with dUTPs. After a final clean up step with a MiniElute Reaction Cleanup kit (Qiagen) and quantification using a Quant-iT Picogreen dsDNA assay in a NanoDrop Fluorospectrometer ND-3300 (Thermo Scientific), the final RNA-seq library preparation (using NEBnext Ultra RNA kit and an UGdase treatment) and directional Illumina sequencing as 100bp paired-end reads was performed at the Vienna BioCenter Core Facilities (VBCF; https://www.viennabiocenter.org/).

We constructed PCA plots (Supplementary Fig. S3) making use of an available *Orchis italica* reference transcriptome (GenBank accession number PRJNA244609; De Paolo *et al*. 2014). The quality-assessed reads were mapped using STAR v.2.6.0c (Dobin et al. 2013)*,* and the 86,080 *Orchis* reference transcripts were then filtered using a count per million minimum threshold of 0.08 across at least eight individuals. This filtering retained 11,424 reference transcripts (13.3%) for further analyses. The expression-based PCA was performed on *RUVs* normalized expression values (as described below) using the *plotPCA* function of the EDAseq v.2.8.0 R package.

We further called SNPs on the aligned RNA-seq reads with freebayes v.1.1.0-46-g8d2b3a0 using a ploidy level of four and ignoring indel and multi-nucleotide polymorphisms (Garrison and Marth 2012). We found 2,709 SNPs that were called across all 19 samples (i.e., with no missing data) distributed across 184 *Orchis italica* transcripts. Other transcripts did not have high enough expression for confident SNP calling in one or more samples. We calculated a genetic relatedness matrix (GRM) using the VanRaden method implemented in snpReady v.0.9.6 R package (VanRaden 2008; Granato et al. 2018).

#### Differential expression analyses

Read trimming was performed with Trimmomatic v.0.33 with LEADING:3 TRAILING:3 SLIDINGWINDOW:4:20 MINLEN:50**.** Trimming quality was then assessed with fastQC*.* The RNA-seq reads were mapped to the *D. incarnata* v.1.0 reference genome with STAR v.2.5.2a (Dobin et al. 2013) The recommended two-pass realignment pipeline was used and reads multi-mapping to more than 20 positions were filtered out. We checked for transcript 5'-3' coverage biases known to affect RNA-seq experiments with RSeQC (Wang et al. 2016). No substantial transcript coverage bias in our data was found (Supplementary Fig. S5). The table of counts was obtained using the *featureCounts* function in Rsubread R package using the full transcripts as features for which to count the reads (Liao et al. 2013).

The *RUVs* normalization approach implemented in the RUVseq v.1.8.0 R package (Risso et al. 2014) was applied to the table of counts for removal of technical variance. *RUVs* uses technical replicates as controls to perform factor analyses on the count matrix and return a linear model that is used to correct for unwanted variance in differential expression analyses. Three (k = 3) factors of unwanted variance were removed as this yielded the best overlap between technical replicates overall; k = 1, 2 and 4 were also tested. We checked these normalizations with Relative Log Expression plots (*plotRLE* function) and Principal Component Analyses plots (*plotPCA* function) using the EDAseq v.2.8.0 R package (Risso et al. 2011). We then fitted the set of read counts using the *glmQLFit* function of *EdgeR* with a non-intercepting model for species and normalizing factors obtained from *RUVs* as fixed effect variables (McCarthy et al. 2012). The *glmQLFTest* function was applied to the fitted model and a $FDR <= 0.05$ threshold for differential expression for the *a priori* species contrast was set.

The genome annotation was then imported into *R* with topGO v.2.26.0 (Alexa and Rahnenfuhrer 2018) and enrichments between pairs of species were done using a two-sided Fisher's exact test against the set of retained transcripts with an FDR smaller than 0.05. Visualization and summary of overrepresented GO terms were performed using the REVIGO web program and results were exported into R. We also used the GOplot v.1.0.2 R package (Supek et al. 2011; Walter et al. 2015). In order to have the directionality of the differential expression towards one species or another and we calculated the z-score for each GO category as $zscore = (\#gene_{up} - \#genes_{down})/\#genes_{total}$.

### Photosynthesis measurements

Over a period of three sunny end-of-May days, we randomized as much as possible the effect of the time of the day by moving between species for each photosynthetic measurement. Because we assumed that our data could be clustered or correlated due to our experimental setup, we fitted the different data obtained from the MultispeQ device with mixed linear models implemented in the lmer function of the lme4 R package (Bates et al. 2015). We fitted a null model with individuals, time of the day and date of the measurements as random effects, and tested the effect of adding species as a fixed variable to the full model. We tested for the significance of species effect on the fitted data using the likelihood ratio test implemented in the anova R function (Bolker et al. 2009).
