## Supplementary Fig. for "Recurrent allopolyploidization events diversify eco-physiological traits in marsh orchids"

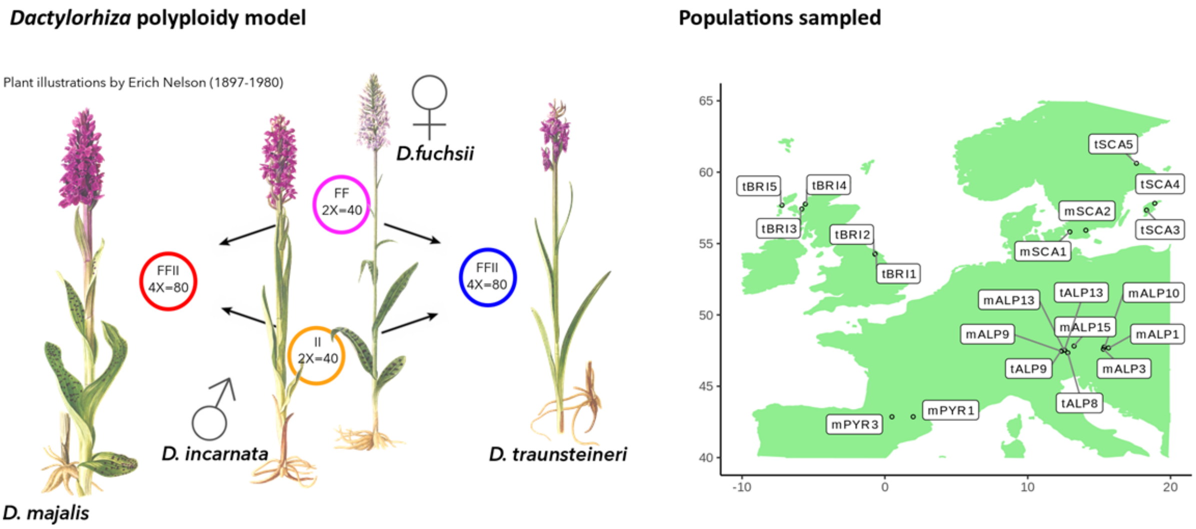


**Figure S1.** Left: The two allotetraploids *D. majalis* and *D. traunsteineri* have both originated from allopolyploidization involving the diploid *D. incarnata* acting as the paternal parent and *D. fuchsii* acting as the maternal parent. The colored bubbles give the chromosome number and genome configuration of each species. Right: Populations sampled for the RNAseq experiment. The acronyms start with a species identifier: m, *D. majalis*; t, *D. traunsteineri*. There are six Alpine populations, four British populations, two populations in the Pyrenees and five Scandinavian populations. Plant illustrations by Erich Nelson (Nelson 1976).

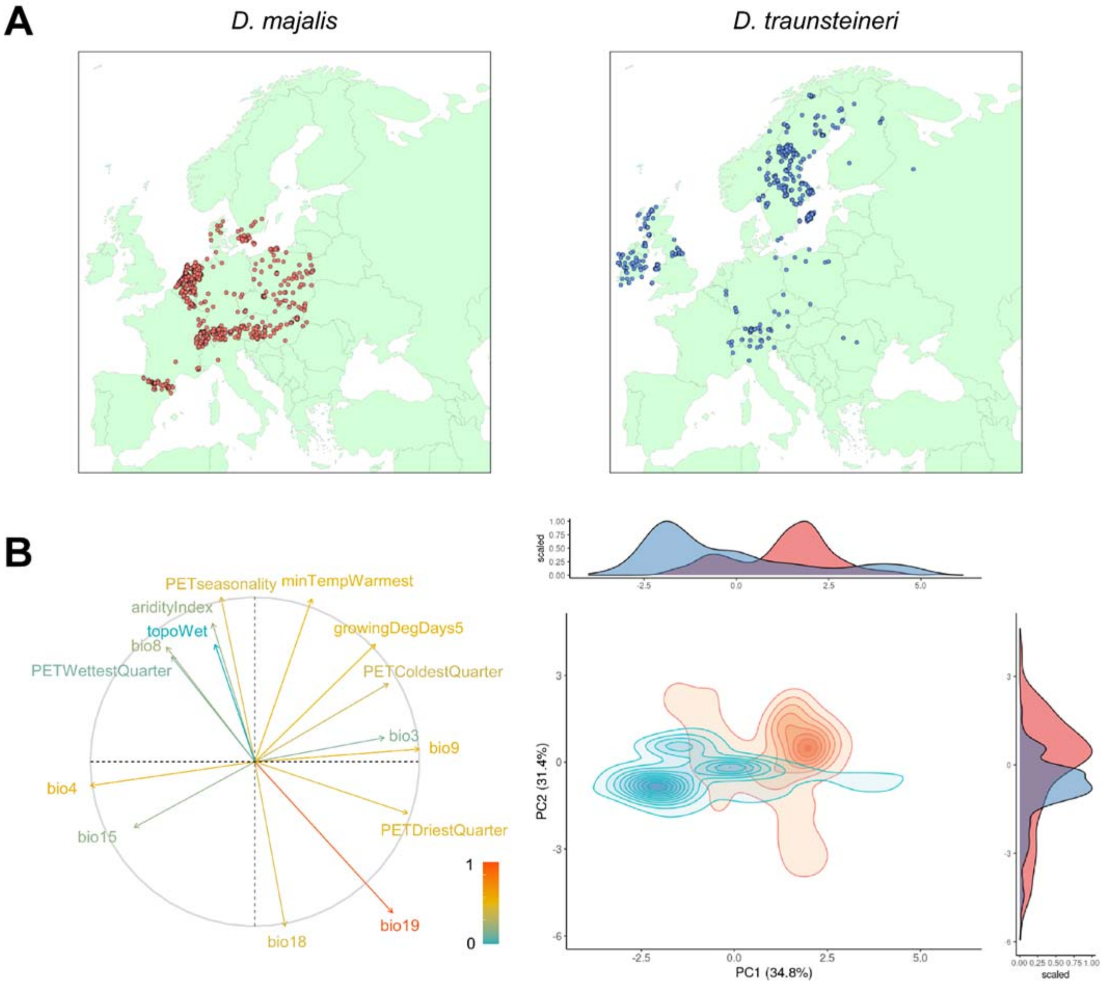


**Figure S2. (A)** Curated *D. majalis* samples (red) and *D. traunsteineri* (blue) extracted from GBIF (accessed February 2018). **(B)** Principal Component Analysis of non-collinear macro-environmental variables performed for GBIF localities and background. The distribution of the selected variables loading on the two main axes is given on the left, with inertia explained by colour of the arrows according to the legend. Bioclim codes are explained in the supplementary text. On the right, density contour plots encompass occurrence points of the *D. majalis* (red) and *D. traunsteineri* (blue) in the 2D environmental space. The occurrence density curves for each axis showed greater environmental divergence in PC1 ($\Delta = 1.60$, $p < 0.001$) compared to PC2 ($\Delta = 0.61$, $p < 0.001$).

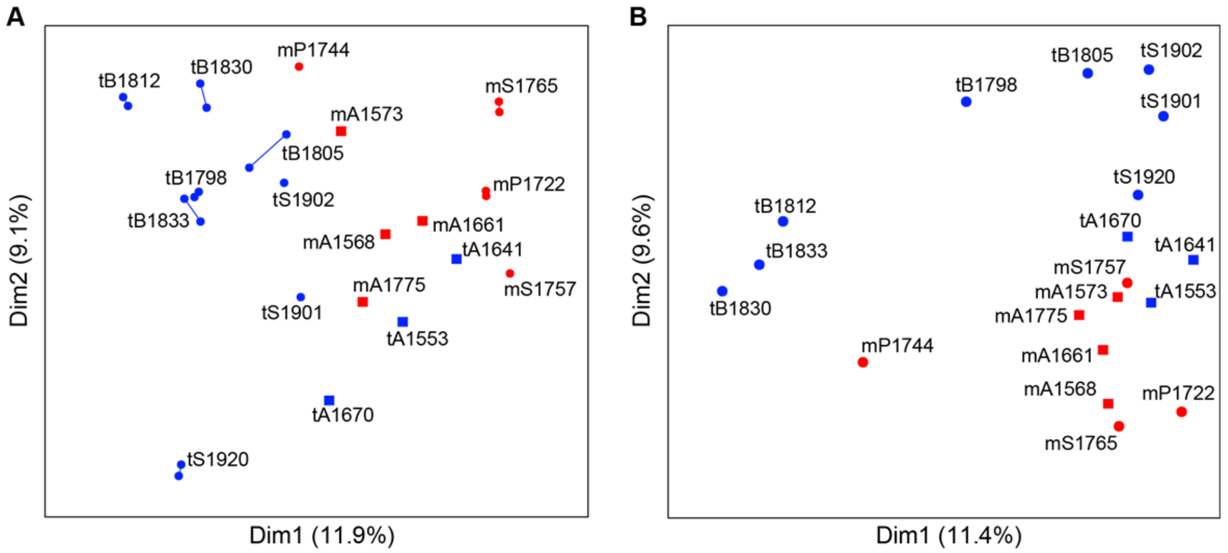


**Figure S3.** Principal component analysis (PCA) for gene expression variation **(A)** and for SNP-based genetic relatedness **(B)** for individuals of *D. majalis* (red symbols) and *D. traunsteineri* (blue) in a common garden setting. Both expression and SNP data show a closer overlap for individuals originating from the sympatric area in the Alps (squares; accession acronym contain "A"), compared with samples from other regions (filled circles; acronyms containing B - British Isles, S - Scandinavia, P - Pyrenees). For panel A, technical replicates are connected with lines.

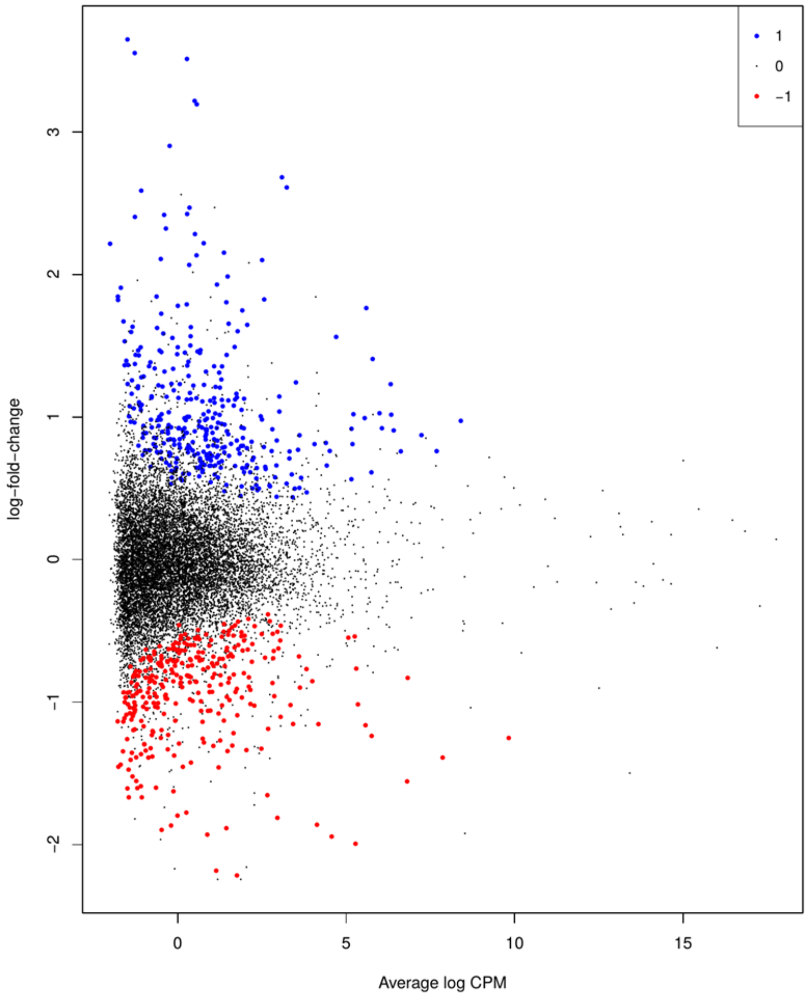


**Figure S4.** Visualization (i.e., MA plot) of the average counts per million on the x-axes and the log fold change on the y-axes (positive values = overexpressed in *D. traunsteineri*; negative values = overexpressed in *D. majalis*). Red points illustrate transcripts that are significantly more highly expressed in *D. majalis* and blue points illustrate transcripts that are more highly expressed in *D. traunsteineri*

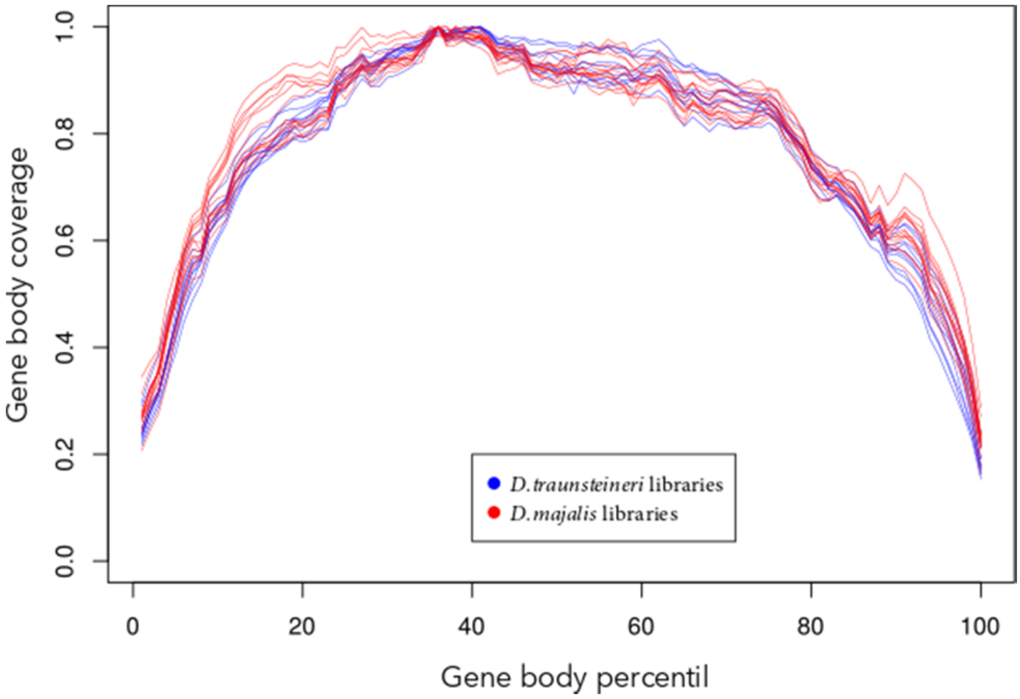


**Figure S5.** Transcriptome gene body coverage, showing that there are no 3' biases in coverage between our different samples. 5'-3' coverage biases are known to affect RNA-seq experiments and were estimated with *RSeQC.*
