## Supplementary Table for "Recurrent allopolyploidization events diversify eco-physiological traits in marsh orchids"

**Table S1.** The populations of *Dactylorhiza majalis* and *D. traunsteineri* we investigated here with different analyses. For leaf chemistry and RNA-seq, the individual accessions are indicated by acronyms including the species (m - *D. majalis*, t - *D. traunsteineri*), the region (A- the Alps, P - the Pyrenees, B - Britain, and S - Scandinavia), and the accession number. Note: the RNA-seq analyses have been performed in a common garden setup; here only the population of origin of the plants is indicated.

| Species | Population ID | Country | Locality | Latitude | Longitude | Soil chemistry | Soil pH | Leaf chemistry | RNA-seq | Photosynthesis |
|---|---|---|---|---|---|---|---|---|---|---|
| majalis | mALP1 | Austria | Altenberg | 47.68553 | 15.6558 | Yes | Yes | mA1031 | mA1568 | - |
| majalis | mALP3 | Austria | Seewiesen | 47.59644 | 15.29366 | - | - | mA1083 | - | - |
| majalis | mALP9 | Austria | Kitzbühl | 47.46155 | 12.3636 | Yes | Yes | mA1651 | - | Yes |
| majalis | mALP10 | Austria | Mooshuben | 47.74485 | 15.35082 | Yes | Yes | mA1478 | mA1573 | - |
| majalis | mALP12 | Austria | Wesenhof | 47.31865 | 11.54238 | - | Yes | - | - | - |
| majalis | mALP13 | Austria | St Ulrich am Pillersee | 47.53075 | 12.57522 | Yes | Yes | mA1667 | mA1661 | Yes |
| majalis | mALP15 | Austria | Fuschlsee | 47.81454 | 13.24461 | Yes | Yes | - | mA1775 | - |
| majalis | mPYR1 | France | D29, Belcaire to Espezel | 42.8617 | 1.980923 | Yes | Yes | mP1003, mP1004 | mP1722 | - |
| majalis | mPYR3 | France | Bourg D'oueil | 42.85863 | 0.495353 | Yes | Yes | mP1744 | mP1744 | - |
| majalis | mSCA1 | Sweden | Lanskrona, Saxtorp | 55.8178 | 12.9456 | Yes | Yes | mS1170 | mS1765 | - |
| majalis | mSCA2 | Sweden | Kristianstadt, Lyngsjön | 55.931 | 14.0683 | Yes | Yes | mS1296 | mS1757 | - |
| traunsteineri | tALP8 | Austria | Prilau | 47.34304 | 12.80421 | Yes | Yes | tA1384, tA1502 | tA1553 | Yes |
| traunsteineri | tALP9 | Austria | Kitzbühl | 47.46098 | 12.36565 | Yes | Yes | tA1429, tA1431 | tA1641 | Yes |
| traunsteineri | tALP11 | Austria | Gaisau | 47.28165 | 11.18457 | - | Yes | - | - | - |
| traunsteineri | tALP13 | Austria | St Ulrich am Pillersee | 47.52927 | 12.57853 | Yes | Yes | tA1627 | tA1670 | Yes |
| traunsteineri | tBRI1 | UK | Sand Dale | 54.25277 | -0.68508 | Yes | Yes | tB1238, tB1239 | tB1798 | - |
| traunsteineri | tBRI2 | UK | Seive Dale Fan | 54.28157 | -0.68977 | - | Yes | tB1806 | tB1805 | - |
| traunsteineri | tBRI3 | UK | Applecross | 57.42202 | -5.81932 | Yes | Yes | tB1180 | tB1812 | - |
| traunsteineri | tBRI4 | UK | Loch Kernsary | 57.767 | -5.569 | - | Yes | tB1811 | - | - |
| traunsteineri | tBRI5 | UK | N Uist, Hebrides | 57.6854 | -7.20566 | Yes | Yes | tB1829 | tB1830, tB1833 | - |
| traunsteineri | tSCA3 | Sweden | Lojsthajd, Gotland | 57.3402 | 18.32118 | Yes | Yes | tS1400 | tS1901, tS1902 | - |
| traunsteineri | tSCA4 | Sweden | Kauparve, Gotland | 57.81707 | 18.89535 | Yes | Yes | tS1416 | - | - |
| traunsteineri | tSCA5 | Sweden | Gårdskär, Uppland | 60.62283 | 17.61094 | Yes | Yes | - | tS1920 | - |

**Table S2.** Results for permutation tests between measurements of various soil chemicals found in the soil for different populations across Europe. Permutation tests were performed because the distributions of the measured chemicals are unknown, we assume each measurement as being independent.

```
########## "N.NO3" ##########

        Exact Permutation Test (network algorithm)

data:  level by species
p-value = 1.149e-06
alternative hypothesis: true mean species=majalis - mean species=traunsteineri is not equal to 0
sample estimates:
mean species=majalis - mean species=traunsteineri
                                         32.65303

########## "N.NH4" ##########

        Permutation Test using Asymptotic Approximation

data:  level by species
Z = -0.37212, p-value = 0.7098
alternative hypothesis: true mean species=majalis - mean species=traunsteineri is not equal to 0
sample estimates:
mean species=majalis - mean species=traunsteineri
                                        -3.500559

########## "Al" ##########

        Permutation Test using Asymptotic Approximation

data:  level by species
Z = 2.212, p-value = 0.02697
alternative hypothesis: true mean species=majalis - mean species=traunsteineri is not equal to 0
sample estimates:
mean species=majalis - mean species=traunsteineri
                                         6.748555

########## "Ca" ##########

        Permutation Test using Asymptotic Approximation

data:  level by species
Z = -1.7507, p-value = 0.08
alternative hypothesis: true mean species=majalis - mean species=traunsteineri is not equal to 0
sample estimates:
mean species=majalis - mean species=traunsteineri
                                        -32.35937

########## "Cd" ##########

        Permutation Test using Asymptotic Approximation

data:  level by species
Z = 1.263, p-value = 0.2066
alternative hypothesis: true mean species=majalis - mean species=traunsteineri is not equal to 0
sample estimates:
mean species=majalis - mean species=traunsteineri
                                        0.1473309

########## "Cr" ##########

        Permutation Test using Asymptotic Approximation

data:  level by species
Z = 2.1042, p-value = 0.03536
alternative hypothesis: true mean species=majalis - mean species=traunsteineri is not equal to 0
sample estimates:
mean species=majalis - mean species=traunsteineri
                                         7.673745

########## "Cu" ##########

        Permutation Test using Asymptotic Approximation

data:  level by species
Z = -0.96389, p-value = 0.3351
alternative hypothesis: true mean species=majalis - mean species=traunsteineri is not equal to 0
sample estimates:
mean species=majalis - mean species=traunsteineri
                                        -16.20464

########## "Fe" ##########

        Permutation Test using Asymptotic Approximation

data:  level by species
Z = 1.1739, p-value = 0.2404
alternative hypothesis: true mean species=majalis - mean species=traunsteineri is not equal to 0
```

**Table S2.** Continued.

```
sample estimates:
mean species=majalis - mean species=traunsteineri
                                         5.935113

########## "K" ##########

        Permutation Test using Asymptotic Approximation

data:  level by species
Z = 3.1316, p-value = 0.001738
alternative hypothesis: true mean species=majalis - mean species=traunsteineri is not equal to 0
sample estimates:
mean species=majalis - mean species=traunsteineri
                                        0.7691066

########## "Mg" ##########

        Permutation Test using Asymptotic Approximation

data:  level by species
Z = -0.57622, p-value = 0.5645
alternative hypothesis: true mean species=majalis - mean species=traunsteineri is not equal to 0
sample estimates:
mean species=majalis - mean species=traunsteineri
                                        -3.772769

########## "Mn" ##########

        Permutation Test using Asymptotic Approximation

data:  level by species
Z = 0.57925, p-value = 0.5624
alternative hypothesis: true mean species=majalis - mean species=traunsteineri is not equal to 0
sample estimates:
mean species=majalis - mean species=traunsteineri
                                         138.7612

########## "Mo" ##########

        Permutation Test using Asymptotic Approximation

data:  level by species
Z = -0.73596, p-value = 0.4618
alternative hypothesis: true mean species=majalis - mean species=traunsteineri is not equal to 0
sample estimates:
mean species=majalis - mean species=traunsteineri
                                       -0.8096167

########## "Na" ##########

        Permutation Test using Asymptotic Approximation

data:  level by species
Z = -0.92668, p-value = 0.3541
alternative hypothesis: true mean species=majalis - mean species=traunsteineri is not equal to 0
sample estimates:
mean species=majalis - mean species=traunsteineri
                                        -55.72212

########## "Ni" ##########

        Permutation Test using Asymptotic Approximation

data:  level by species
Z = 2.212, p-value = 0.02696
alternative hypothesis: true mean species=majalis - mean species=traunsteineri is not equal to 0
sample estimates:
mean species=majalis - mean species=traunsteineri
                                         9.028801

########## "P" ##########

        Permutation Test using Asymptotic Approximation

data:  level by species
Z = 3.4124, p-value = 0.000644
alternative hypothesis: true mean species=majalis - mean species=traunsteineri is not equal to 0
sample estimates:
mean species=majalis - mean species=traunsteineri
                                         416.8851

########## "Pb" ##########

        Permutation Test using Asymptotic Approximation

data:  level by species
Z = 0.7782, p-value = 0.4365
```

**Table S2.** Continued.

```
alternative hypothesis: true mean species=majalis - mean species=traunsteineri is not equal to 0
sample estimates:
mean species=majalis - mean species=traunsteineri
                                         6.103012

########## "S" ##########

        Permutation Test using Asymptotic Approximation

data:  level by species
Z = 1.0853, p-value = 0.2778
alternative hypothesis: true mean species=majalis - mean species=traunsteineri is not equal to 0
sample estimates:
mean species=majalis - mean species=traunsteineri
                                         703.0451

########## "Zn" ##########

        Permutation Test using Asymptotic Approximation

data:  level by species
Z = 0.80442, p-value = 0.4212
alternative hypothesis: true mean species=majalis - mean species=traunsteineri is not equal to 0
sample estimates:
mean species=majalis - mean species=traunsteineri
                                         17.15471

########## "pH" ##########

        Permutation Test using Asymptotic Approximation

data:  trapH$pH and majpH$pH
Z = 4.816, p-value = 1.464e-06
alternative hypothesis: true mean trapH$pH - mean majpH$pH is not equal to 0
sample estimates:
mean trapH$pH - mean majpH$pH
                    0.7143187
```

**Table S3.** Table showing enrichment GO terms with their corresponding name. The third column is the total number of annotated transcripts in the corresponding category; the fourth column is the number of significant transcripts differentially expressed between *D. traunsteineri* and *D. majalis*; the fifth column is the expected number of significant transcripts that should be differentially expressed between both species under a null model; the sixth column is the p-value for the corresponding GO enrichment, and the seventh column is the corresponding GO branch. BP is Biological Process, MF is Molecular Function, and CC is Cellular Component.

| GO.ID | Term | Annotated | Significant | Expected | weight01_pval | branch |
|---|---|---|---|---|---|---|
| GO:0009768 | photosynthesis, light harvesting in phot... | 15 | 8 | 0.85 | 0.00000046 | BP |
| GO:0071555 | cell wall organization | 173 | 26 | 9.83 | 0.0000079 | BP |
| GO:0009269 | response to desiccation | 6 | 4 | 0.34 | 0.00014 | BP |
| GO:0015995 | chlorophyll biosynthetic process | 45 | 9 | 2.56 | 0.00032 | BP |
| GO:0015979 | photosynthesis | 204 | 36 | 11.59 | 0.00092 | BP |
| GO:0010411 | xyloglucan metabolic process | 19 | 6 | 1.08 | 0.00149 | BP |
| GO:0006809 | nitric oxide biosynthetic process | 10 | 4 | 0.57 | 0.00164 | BP |
| GO:0010207 | photosystem II assembly | 11 | 4 | 0.62 | 0.00246 | BP |
| GO:0045454 | cell redox homeostasis | 75 | 11 | 4.26 | 0.00318 | BP |
| GO:0009769 | photosynthesis, light harvesting in phot... | 2 | 2 | 0.11 | 0.00322 | BP |
| GO:0071435 | potassium ion export | 2 | 2 | 0.11 | 0.00322 | BP |
| GO:1902265 | abscisic acid homeostasis | 2 | 2 | 0.11 | 0.00322 | BP |
| GO:0018298 | protein-chromophore linkage | 67 | 10 | 3.81 | 0.00424 | BP |
| GO:0006662 | glycerol ether metabolic process | 21 | 5 | 1.19 | 0.00551 | BP |
| GO:0009767 | photosynthetic electron transport chain | 40 | 10 | 2.27 | 0.00629 | BP |
| GO:0045493 | xylan catabolic process | 8 | 3 | 0.45 | 0.00823 | BP |
| GO:0009637 | response to blue light | 40 | 7 | 2.27 | 0.00828 | BP |
| GO:0031022 | nuclear migration along microfilament | 3 | 2 | 0.17 | 0.00929 | BP |
| GO:0010345 | suberin biosynthetic process | 9 | 3 | 0.51 | 0.01182 | BP |
| GO:0070413 | trehalose metabolism in response to stre... | 9 | 3 | 0.51 | 0.01182 | BP |
| GO:0010193 | response to ozone | 17 | 4 | 0.97 | 0.01353 | BP |
| GO:0006782 | protoporphyrinogen IX biosynthetic proce... | 10 | 3 | 0.57 | 0.01619 | BP |
| GO:0010206 | photosystem II repair | 10 | 3 | 0.57 | 0.01619 | BP |
| GO:0010119 | regulation of stomatal movement | 30 | 6 | 1.7 | 0.01647 | BP |
| GO:0042546 | cell wall biogenesis | 82 | 8 | 4.66 | 0.01659 | BP |
| GO:2000785 | regulation of autophagosome assembly | 4 | 2 | 0.23 | 0.01789 | BP |
| GO:0009416 | response to light stimulus | 348 | 33 | 19.77 | 0.01971 | BP |
| GO:0005992 | trehalose biosynthetic process | 11 | 3 | 0.62 | 0.02133 | BP |
| GO:1901002 | positive regulation of response to salt ... | 11 | 3 | 0.62 | 0.02133 | BP |
| GO:0009773 | photosynthetic electron transport in pho... | 12 | 3 | 0.68 | 0.02727 | BP |
| GO:0051103 | DNA ligation involved in DNA repair | 5 | 2 | 0.28 | 0.0287 | BP |
| GO:0070588 | calcium ion transmembrane transport | 5 | 2 | 0.28 | 0.0287 | BP |
| GO:0009816 | defense response to bacterium, incompati... | 25 | 5 | 1.42 | 0.03307 | BP |
| GO:0009409 | response to cold | 148 | 14 | 8.41 | 0.03467 | BP |
| GO:0009753 | response to jasmonic acid | 90 | 10 | 5.11 | 0.0363 | BP |
| GO:0009651 | response to salt stress | 229 | 21 | 13.01 | 0.03705 | BP |
| GO:0009835 | fruit ripening | 6 | 2 | 0.34 | 0.04146 | BP |
| GO:0010224 | response to UV-B | 35 | 5 | 1.99 | 0.04596 | BP |
| GO:0055114 | oxidation-reduction process | 616 | 50 | 34.99 | 0.04817 | BP |
| GO:0009695 | jasmonic acid biosynthetic process | 15 | 3 | 0.85 | 0.04973 | BP |
| GO:0031409 | pigment binding | 14 | 7 | 0.75 | 0.0000031 | MF |
| GO:0008559 | xenobiotic-transporting ATPase activity | 15 | 7 | 0.81 | 0.0000055 | MF |
| GO:0016168 | chlorophyll binding | 50 | 10 | 2.69 | 0.00027 | MF |
| GO:0016762 | xyloglucan:xyloglucosyl transferase acti... | 9 | 4 | 0.48 | 0.00084 | MF |
| GO:0003825 | alpha,alpha-trehalose-phosphate synthase... | 6 | 3 | 0.32 | 0.00274 | MF |
| GO:0003950 | NAD+ ADP-ribosyltransferase activity | 6 | 3 | 0.32 | 0.00274 | MF |
| GO:0004805 | trehalose-phosphatase activity | 6 | 3 | 0.32 | 0.00274 | MF |
| GO:0009044 | xylan 1,4-beta-xylosidase activity | 6 | 3 | 0.32 | 0.00274 | MF |
| GO:0050464 | nitrate reductase (NADPH) activity | 2 | 2 | 0.11 | 0.00289 | MF |
| GO:0019904 | protein domain specific binding | 49 | 9 | 2.64 | 0.00366 | MF |
| GO:0004565 | beta-galactosidase activity | 14 | 4 | 0.75 | 0.00537 | MF |

**Table S3.** Continued

| GO.ID | Term | Annotated | Significant | Expected | weight01_pval | branch |
|---|---|---|---|---|---|---|
| GO:0009703 | nitrate reductase (NADH) activity | 3 | 2 | 0.16 | 0.00835 | MF |
| GO:0046556 | alpha-L-arabinofuranosidase activity | 3 | 2 | 0.16 | 0.00835 | MF |
| GO:0043546 | molybdopterin cofactor binding | 3 | 2 | 0.16 | 0.00835 | MF |
| GO:0004497 | monooxygenase activity | 80 | 11 | 4.3 | 0.01278 | MF |
| GO:0015035 | protein disulfide oxidoreductase activit... | 38 | 6 | 2.04 | 0.01492 | MF |
| GO:0004742 | dihydrolipoyllysine-residue acetyltransf... | 4 | 2 | 0.22 | 0.01611 | MF |
| GO:0005544 | calcium-dependent phospholipid binding | 4 | 2 | 0.22 | 0.01611 | MF |
| GO:0016984 | ribulose-bisphosphate carboxylase activi... | 4 | 2 | 0.22 | 0.01611 | MF |
| GO:0001653 | peptide receptor activity | 4 | 2 | 0.22 | 0.01611 | MF |
| GO:0016874 | ligase activity | 152 | 13 | 8.18 | 0.01769 | MF |
| GO:0020037 | heme binding | 100 | 11 | 5.38 | 0.01812 | MF |
| GO:0003700 | transcription factor activity, sequence-... | 399 | 31 | 21.46 | 0.01935 | MF |
| GO:0046910 | pectinesterase inhibitor activity | 12 | 3 | 0.65 | 0.02364 | MF |
| GO:0004791 | thioredoxin-disulfide reductase activity | 21 | 4 | 1.13 | 0.02383 | MF |
| GO:0008061 | chitin binding | 5 | 2 | 0.27 | 0.0259 | MF |
| GO:0003910 | DNA ligase (ATP) activity | 5 | 2 | 0.27 | 0.0259 | MF |
| GO:0010277 | chlorophyllide a oxygenase [overall] act... | 5 | 2 | 0.27 | 0.0259 | MF |
| GO:0050660 | flavin adenine dinucleotide binding | 56 | 7 | 3.01 | 0.02648 | MF |
| GO:0047134 | protein-disulfide reductase activity | 22 | 4 | 1.18 | 0.02793 | MF |
| GO:0016705 | oxidoreductase activity, acting on paire... | 99 | 10 | 5.33 | 0.02886 | MF |
| GO:0001053 | plastid sigma factor activity | 6 | 2 | 0.32 | 0.03748 | MF |
| GO:0004674 | protein serine/threonine kinase activity | 452 | 32 | 24.31 | 0.04355 | MF |
| GO:0030246 | carbohydrate binding | 112 | 10 | 6.02 | 0.04437 | MF |
| GO:0009535 | chloroplast thylakoid membrane | 304 | 47 | 16.48 | 0.000000017 | CC |
| GO:0005618 | cell wall | 180 | 24 | 9.76 | 0.0000024 | CC |
| GO:0048046 | apoplast | 106 | 19 | 5.75 | 0.0000034 | CC |
| GO:0010287 | plastoglobule | 47 | 12 | 2.55 | 0.0000051 | CC |
| GO:0009522 | photosystem I | 43 | 16 | 2.33 | 0.000013 | CC |
| GO:0009538 | photosystem I reaction center | 5 | 4 | 0.27 | 0.000041 | CC |
| GO:0009523 | photosystem II | 62 | 16 | 3.36 | 0.00013 | CC |
| GO:0009941 | chloroplast envelope | 351 | 33 | 19.03 | 0.00014 | CC |
| GO:0000325 | plant-type vacuole | 79 | 9 | 4.28 | 0.00021 | CC |
| GO:0009654 | photosystem II oxygen evolving complex | 17 | 5 | 0.92 | 0.00164 | CC |
| GO:0000795 | synaptonemal complex | 6 | 3 | 0.33 | 0.0028 | CC |
| GO:0009341 | beta-galactosidase complex | 6 | 3 | 0.33 | 0.0028 | CC |
| GO:0098807 | chloroplast thylakoid membrane protein c... | 6 | 6 | 0.33 | 0.00288 | CC |
| GO:0009517 | PSII associated light-harvesting complex... | 2 | 2 | 0.11 | 0.00293 | CC |
| GO:0030093 | chloroplast photosystem I | 2 | 2 | 0.11 | 0.00293 | CC |
| GO:0005576 | extracellular region | 350 | 39 | 18.97 | 0.00369 | CC |
| GO:0009543 | chloroplast thylakoid lumen | 40 | 7 | 2.17 | 0.00515 | CC |
| GO:0016021 | integral component of membrane | 2002 | 128 | 108.52 | 0.00868 | CC |
| GO:0019898 | extrinsic component of membrane | 51 | 8 | 2.76 | 0.01936 | CC |
| GO:0009574 | preprophase band | 5 | 2 | 0.27 | 0.02628 | CC |
| GO:0009579 | thylakoid | 389 | 58 | 21.09 | 0.02932 | CC |
| GO:0009570 | chloroplast stroma | 389 | 40 | 21.09 | 0.02969 | CC |

**Table S4.** Linear mixed model fitting and significance results for various measurements obtained with MicroSpec v-1.0. Photosynthesis measurements were fitted with individuals, time, and date of the measurements modeled as random effects and species modeled as a fixed variable.  All measurements were done for populations growing in the Alps in an interval of three days.

```
########## "Ambient_Humidity" ##########
Data: dtFspecMT
Models:
lmm.null: get(d) ~ 1 + (1 | Time_of_Day) + (1 | sample_name) + (1 | date)
lmm.full: get(d) ~ species + (1 | Time_of_Day) + (1 | sample_name) + (1 |
lmm.full:     date)
         Df    AIC    BIC  logLik deviance  Chisq Chi Df Pr(>Chisq)
lmm.null  5 422,30 434,14 -206,15   412,30
lmm.full  6 416,25 430,46 -202,12   404,25 8,0484      1   0,004554 **
---
Signif. codes:  0    ***    0,001    **    0,01    *    0,05    .    0,1        1
---------- SUMMARY ----------
Linear mixed model fit by maximum likelihood  ['lmerMod']
Formula: get(d) ~ species + (1 | Time_of_Day) + (1 | sample_name) + (1 |      date)
   Data: dtFspecMT

     AIC      BIC   logLik deviance df.resid
   416.2    430.5   -202.1    404.2       73

Scaled residuals:
     Min       1Q   Median       3Q      Max
-2.13244 -0.34430 -0.00416  0.33198  2.87213

Random effects:
 Groups      Name        Variance Std.Dev.
 sample_name (Intercept)  8.040   2.835
 Time_of_Day (Intercept)  5.356   2.314
 date        (Intercept) 53.107   7.287
 Residual                 2.496   1.580
Number of obs: 79, groups:  sample_name, 46; Time_of_Day, 7; date, 4

Fixed effects:
                     Estimate Std. Error t value
(Intercept)            56.181      3.862  14.546
speciestraunsteineri    3.467      1.129   3.072

Correlation of Fixed Effects:
            (Intr)
spcstrnstnr -0.188

########## "Ambient_Temperature" ##########
Data: dtFspecMT
Models:
lmm.null: get(d) ~ 1 + (1 | Time_of_Day) + (1 | sample_name) + (1 | date)
lmm.full: get(d) ~ species + (1 | Time_of_Day) + (1 | sample_name) + (1 |
lmm.full:     date)
         Df    AIC    BIC  logLik deviance  Chisq Chi Df Pr(>Chisq)
lmm.null  5 279,93 291,78 -134,97   269,93
lmm.full  6 241,47 255,68 -114,73   229,47 40,468      1  1,999e-10 ***
---
Signif. codes:  0    ***    0,001    **    0,01    *    0,05    .    0,1        1
---------- SUMMARY ----------
Linear mixed model fit by maximum likelihood  ['lmerMod']
Formula: get(d) ~ species + (1 | Time_of_Day) + (1 | sample_name) + (1 |      date)
   Data: dtFspecMT

     AIC      BIC   logLik deviance df.resid
   241.5    255.7   -114.7    229.5       73

Scaled residuals:
     Min       1Q   Median       3Q      Max
-2.18307 -0.34817  0.00108  0.32716  1.96516

Random effects:
 Groups      Name        Variance Std.Dev.
 sample_name (Intercept)  1.0920  1.0450
 Time_of_Day (Intercept) 14.6324  3.8252
 date        (Intercept)  3.8794  1.9696
 Residual                 0.1486  0.3855
Number of obs: 79, groups:  sample_name, 46; Time_of_Day, 7; date, 4

Fixed effects:
                     Estimate Std. Error t value
(Intercept)           25.9381     1.7891  14.497
speciestraunsteineri   4.0771     0.4406   9.253

Correlation of Fixed Effects:
            (Intr)
spcstrnstnr -0.159

########## "Leaf_Angle" ##########
Data: dtFspecMT
Models:
lmm.null: get(d) ~ 1 + (1 | Time_of_Day) + (1 | sample_name) + (1 | date)
```

**Table S4.** Continued

```
lmm.full: get(d) ~ species + (1 | Time_of_Day) + (1 | sample_name) + (1 |
lmm.full:     date)
         Df    AIC    BIC  logLik deviance  Chisq Chi Df Pr(>Chisq)
lmm.null  5 700,37 712,22 -345,19   690,37
lmm.full  6 702,31 716,53 -345,16   690,31 0,0626      1     0,8024

########## "Leaf.Temp.Differential" ##########
Data: dtFspecMT
Models:
lmm.null: get(d) ~ 1 + (1 | Time_of_Day) + (1 | sample_name) + (1 | date)
lmm.full: get(d) ~ species + (1 | Time_of_Day) + (1 | sample_name) + (1 |
lmm.full:     date)
         Df    AIC    BIC  logLik deviance  Chisq Chi Df Pr(>Chisq)
lmm.null  5 205,01 216,86 -97,506   195,01
lmm.full  6 206,94 221,16 -97,472   194,94 0,0679      1     0,7944

########## "LEF" ##########
Data: dtFspecMT
Models:
lmm.null: get(d) ~ 1 + (1 | Time_of_Day) + (1 | sample_name) + (1 | date)
lmm.full: get(d) ~ species + (1 | Time_of_Day) + (1 | sample_name) + (1 |
lmm.full:     date)
         Df    AIC    BIC  logLik deviance  Chisq Chi Df Pr(>Chisq)
lmm.null  5 682,50 694,35 -336,25   672,50
lmm.full  6 681,45 695,67 -334,73   669,45 3,0467      1     0,0809 .
---
Signif. codes:  0    ***    0,001    **    0,01    *    0,05    .    0,1         1

########## "PAR" ##########
Data: dtFspecMT
Models:
lmm.null: get(d) ~ 1 + (1 | Time_of_Day) + (1 | sample_name) + (1 | date)
lmm.full: get(d) ~ species + (1 | Time_of_Day) + (1 | sample_name) + (1 |
lmm.full:     date)
         Df    AIC    BIC  logLik deviance  Chisq Chi Df Pr(>Chisq)
lmm.null  5 987,27 999,11 -488,63   977,27
lmm.full  6 985,10 999,32 -486,55   973,10 4,1642      1    0,04129 *
---
Signif. codes:  0    ***    0,001    **    0,01    *    0,05    .    0,1         1
---------- SUMMARY ----------
Linear mixed model fit by maximum likelihood  ['lmerMod']
Formula: get(d) ~ species + (1 | Time_of_Day) + (1 | sample_name) + (1 |      date)
   Data: dtFspecMT

     AIC      BIC   logLik deviance df.resid
   985.1    999.3   -486.6    973.1       73

Scaled residuals:
    Min      1Q  Median      3Q     Max
-1.8419 -0.4497 -0.1083  0.3832  1.9450

Random effects:
 Groups      Name        Variance Std.Dev.
 sample_name (Intercept) 10062    100.31
 Time_of_Day (Intercept)  2080     45.61
 date        (Intercept)  1241     35.23
 Residual                 5200     72.11
Number of obs: 79, groups:  sample_name, 46; Time_of_Day, 7; date, 4

Fixed effects:
                     Estimate Std. Error t value
(Intercept)            244.71      39.75   6.156
speciestraunsteineri    82.00      38.66   2.121

Correlation of Fixed Effects:
            (Intr)
spcstrnstnr -0.624

########## "NPQt" ##########
Data: dtFspecMT
Models:
lmm.null: get(d) ~ 1 + (1 | Time_of_Day) + (1 | sample_name) + (1 | date)
lmm.full: get(d) ~ species + (1 | Time_of_Day) + (1 | sample_name) + (1 |
lmm.full:     date)
         Df    AIC    BIC  logLik deviance  Chisq Chi Df Pr(>Chisq)
lmm.null  5 84,801 96,649 -37,401   74,801
lmm.full  6 80,908 95,124 -34,454   68,908 5,8937      1    0,01519 *
---
Signif. codes:  0    ***    0,001    **    0,01    *    0,05    .    0,1         1
---------- SUMMARY ----------
Linear mixed model fit by maximum likelihood  ['lmerMod']
Formula: get(d) ~ species + (1 | Time_of_Day) + (1 | sample_name) + (1 |      date)
   Data: dtFspecMT

     AIC      BIC   logLik deviance df.resid
    80.9     95.1    -34.5     68.9       73
```

**Table S4.** Continued

```
Scaled residuals:
    Min      1Q  Median      3Q     Max
-1.4270 -0.4594 -0.1077  0.3824  1.9979

Random effects:
 Groups      Name        Variance Std.Dev.
 sample_name (Intercept) 0.1495   0.3867
 Time_of_Day (Intercept) 0.0000   0.0000
 date        (Intercept) 0.1017   0.3188
 Residual                0.0404   0.2010
Number of obs: 79, groups:  sample_name, 46; Time_of_Day, 7; date, 4

Fixed effects:
                     Estimate Std. Error t value
(Intercept)            0.7280     0.1948   3.737
speciestraunsteineri   0.3428     0.1327   2.584

Correlation of Fixed Effects:
            (Intr)
spcstrnstnr -0.460

########## "Phi2" ##########
Data: dtFspecMT
Models:
lmm.null: get(d) ~ 1 + (1 | Time_of_Day) + (1 | sample_name) + (1 | date)
lmm.full: get(d) ~ species + (1 | Time_of_Day) + (1 | sample_name) + (1 |
lmm.full:     date)
         Df     AIC     BIC logLik deviance  Chisq Chi Df Pr(>Chisq)
lmm.null  5 -224,16 -212,31 117,08  -234,16
lmm.full  6 -231,92 -217,70 121,96  -243,92 9,7598      1   0,001784 **
---
Signif. codes:  0    ***    0,001    **    0,01    *    0,05    .    0,1         1
---------- SUMMARY ----------
Linear mixed model fit by maximum likelihood  ['lmerMod']
Formula: get(d) ~ species + (1 | Time_of_Day) + (1 | sample_name) + (1 |      date)
   Data: dtFspecMT

     AIC      BIC   logLik deviance df.resid
  -231.9   -217.7    122.0   -243.9       73

Scaled residuals:
    Min      1Q  Median      3Q     Max
-2.0848 -0.4408 -0.0179  0.5920  1.4665

Random effects:
 Groups      Name        Variance  Std.Dev.
 sample_name (Intercept) 2.152e-03 4.639e-02
 Time_of_Day (Intercept) 4.577e-04 2.139e-02
 date        (Intercept) 2.084e-18 1.444e-09
 Residual                1.065e-03 3.263e-02
Number of obs: 79, groups:  sample_name, 46; Time_of_Day, 7; date, 4

Fixed effects:
                     Estimate Std. Error t value
(Intercept)           0.48411    0.01618   29.93
speciestraunsteineri -0.05933    0.01771   -3.35

Correlation of Fixed Effects:
            (Intr)
spcstrnstnr -0.694

########## "PhiNO" ##########
Data: dtFspecMT
Models:
lmm.null: get(d) ~ 1 + (1 | Time_of_Day) + (1 | sample_name) + (1 | date)
lmm.full: get(d) ~ species + (1 | Time_of_Day) + (1 | sample_name) + (1 |
lmm.full:     date)
         Df     AIC     BIC logLik deviance  Chisq Chi Df Pr(>Chisq)
lmm.null  5 -236,00 -224,15 123,00  -246,00
lmm.full  6 -236,27 -222,05 124,13  -248,27 2,2715      1     0,1318

########## "PhiNPQ" ##########
Data: dtFspecMT
Models:
lmm.null: get(d) ~ 1 + (1 | Time_of_Day) + (1 | sample_name) + (1 | date)
lmm.full: get(d) ~ species + (1 | Time_of_Day) + (1 | sample_name) + (1 |
lmm.full:     date)
         Df     AIC     BIC logLik deviance  Chisq Chi Df Pr(>Chisq)
lmm.null  5 -198,11 -186,26 104,05  -208,11
lmm.full  6 -204,95 -190,74 108,48  -216,95 8,8481      1   0,002934 **
---
Signif. codes:  0    ***    0,001    **    0,01    *    0,05    .    0,1         1
---------- SUMMARY ----------
Linear mixed model fit by maximum likelihood  ['lmerMod']
Formula: get(d) ~ species + (1 | Time_of_Day) + (1 | sample_name) + (1 |      date)
   Data: dtFspecMT
```

**Table S4.** Continued

```
     AIC      BIC   logLik deviance df.resid
  -205.0   -190.7    108.5   -217.0       73

Scaled residuals:
    Min      1Q  Median      3Q     Max
-2.2984 -0.4329 -0.1064  0.4454  2.1185

Random effects:
 Groups      Name        Variance  Std.Dev.
 sample_name (Intercept) 0.0038732 0.06223
 Time_of_Day (Intercept) 0.0009791 0.03129
 date        (Intercept) 0.0008445 0.02906
 Residual                0.0010926 0.03305
Number of obs: 79, groups:  sample_name, 46; Time_of_Day, 7; date, 4

Fixed effects:
                     Estimate Std. Error t value
(Intercept)           0.21119    0.02647   7.977
speciestraunsteineri  0.07903    0.02302   3.433

Correlation of Fixed Effects:
            (Intr)
spcstrnstnr -0.559

########## "Relative_Chlorophyll" ##########
Data: dtFspecMT
Models:
lmm.null: get(d) ~ 1 + (1 | Time_of_Day) + (1 | sample_name) + (1 | date)
lmm.full: get(d) ~ species + (1 | Time_of_Day) + (1 | sample_name) + (1 |
lmm.full:     date)
         Df    AIC    BIC  logLik deviance  Chisq Chi Df Pr(>Chisq)
lmm.null  5 541,90 553,75 -265,95   531,90
lmm.full  6 537,79 552,01 -262,90   525,79 6,1055      1    0,01348 *
---
Signif. codes:  0    ***    0,001    **    0,01    *    0,05    .    0,1         1
---------- SUMMARY ----------
Linear mixed model fit by maximum likelihood  ['lmerMod']
Formula: get(d) ~ species + (1 | Time_of_Day) + (1 | sample_name) + (1 |      date)
   Data: dtFspecMT

     AIC      BIC   logLik deviance df.resid
   537.8    552.0   -262.9    525.8       73

Scaled residuals:
     Min       1Q   Median       3Q      Max
-1.92730 -0.43289 -0.03472  0.46551  2.17496

Random effects:
 Groups      Name        Variance Std.Dev.
 sample_name (Intercept) 31.2130  5.5869
 Time_of_Day (Intercept)  0.0000  0.0000
 date        (Intercept)  0.1999  0.4471
 Residual                22.6397  4.7581
Number of obs: 79, groups:  sample_name, 46; Time_of_Day, 7; date, 4

Fixed effects:
                     Estimate Std. Error t value
(Intercept)            42.931      1.646  26.088
speciestraunsteineri   -5.327      2.053  -2.595

Correlation of Fixed Effects:
            (Intr)
spcstrnstnr -0.784
```
